# High-Resolution Magnetization-Transfer Imaging of *Post-Mortem* Marmoset Brain: Comparisons with Relaxometry and Histology

**DOI:** 10.1101/2022.09.05.506600

**Authors:** Henrik Marschner, André Pampel, Roland Müller, Katja Reimann, Nicolas Bock, Markus Morawski, Stefan Geyer, Harald E. Möller

**Affiliations:** Max Planck Institute for Human Cognitive and Brain Sciences, Stephanstraße 1a, 04103 Leipzig, Germany; Leipzig University, Paul Flechsig Institute of Brain Research, Liebigstraße 19, 04103 Leipzig, Germany; McMaster University, Department of Psychology, Neuroscience & Behaviour, 1280 Main Street West, Hamilton, ON L8S4K1, Canada

**Keywords:** Histology, iron, longitudinal relaxation, macromolecular pool, magnetization transfer, myelin, *T*_1_, 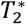, transverse relaxation

## Abstract

Cell membranes and macromolecules or paramagnetic compounds interact with water proton spins, which modulates magnetic resonance imaging (MRI) contrast providing information on tissue composition. For a further investigation, quantitative magnetization transfer (qMT) parameters (at 3T), including the ratio of the macromolecular and water proton pools, ℱ, and the exchange-rate constant as well as the (observed) longitudinal and the effective transverse relaxation rates (at 3T and 7T), 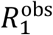 and 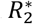 respectively, were measured at high spatial resolution (200 μm) in a slice of fixed marmoset brain and compared to histology results obtained with Gallyas’ myelin stain and Perls’ iron stain. 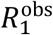 and 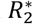 were linearly correlated with the iron content for the entire slice, whereas distinct differences were obtained between gray and white matter for correlations of relaxometry and qMT parameters with myelin content. The combined results suggest that the macromolecular pool interacting with water consists of myelin and (less efficient) non-myelin contributions. Despite strong correlation of ℱ and 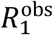 none of these parameters was uniquely specific to myelination. Due to additional sensitivity to iron stores, 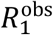 and 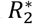 were more sensitive for depicting microstructural differences between cortical layers than ℱ.

**Highlights:**

- MRI (200μm) is correlated with myelin and iron histology in fixed marmoset brain.
- Detailed *z*-spectra are employed for precise magnetization-transfer (MT) measurements.
- Longitudinal and effective transverse relaxation rates depend linearly on tissue iron.
- Longitudinal relaxation and MT are not uniquely specific to myelin.
- Myelin and non-myelin macromolecules impact water relaxation and MT contrast.

## 1 Introduction

Quantitative magnetization transfer imaging (qMTI) is a versatile tool to obtain information on cell membranes or other macromolecular compounds (*e.g*., proteins) via cross relaxation or proton exchange with water molecules (Edzes & Samulski, 1977, 1978; Sled, 2018; Wolff & Balaban, 1989). Such semisolid components are not directly visible to standard magnetic resonance imaging (MRI) due to their very short transverse relaxation time, *T*_2_. In a typical MT experiment, the semisolid spin pool is saturated by radiofrequency (RF) irradiation of limited bandwidth applied off resonance of the narrow water line. This leads to a transient water-signal change that can be fitted to a set of differential equations, such as the binary spin-bath (BSB) model (Henkelman *et al*., 1993). Thereby, information is obtained on the semisolid pool and magnetization exchange rates.

The efficiency of the magnetization transfer (MT) depends on the presence and number of binding sites for water on the semisolid components and on the dynamics of the system (Bryant & Korb, 2005). In brain tissue, and especially in white matter (WM), the most important contribution to cross-relaxation results from myelin that envelops the axons (Koenig *et al*., 1990; Laule *et al*., 2007; Möller *et al*., 2019; Sled, 2018). In particular, galactolipids and cholesterol in the myelin membrane have been proposed as sites with efficient coupling to water molecules (Ceckler *et al*., 1992; Fralix *et al*., 1991; Koenig, 1991, Kucharczyk *et al*., 1994). Previous work has also demonstrated correlations between the relative size of the semisolid pool estimated by qMTI in selected regions of interest (ROIs) and histological measures of myelin content (Schmierer *et al*., 2007).

The goal of the current work was a more comprehensive, voxel-by-voxel comparison of qMTI of fixed brain and histology. To put the results into a broader context, measurements of the longitudinal rate *R*_1_= 1/*T*_1_ and the effective transverse relaxation rate 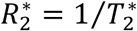 were also integrated in the experiments. Recently, there has been growing interest in using the common marmoset (*Callithrix jacchus*), a New World monkey, in neuroimaging research (Bock *et al*., 2009; Liu *et al*., 2011; Newman *et al*., 2009). While it demonstrates typical primate brain functional organization, its small, lissencephalic brain has no complex folding pattern offering excellent conditions for cortical imaging at high spatial resolution. Further, the brain’s overall gray-to-white matter ratio is much higher than in rodent imaging models, making the marmoset an ideal for quantitative studies of myelin in deep brain structures.

## 2 Experimental Procedures

### 2.1 Brain Specimen

The animal procedures were approved by the NINDS Animal Care and Use Committee. The brain of a male common marmoset (*Callithrix jacchus*) that had died from natural causes at an age of 7.6 years was dissected out and fixed by perfusion with 4% formalin in phosphate-buffered saline (PBS). Subsequently, the specimen was stored in PBS with 0.1% sodium azide (NaN_3_) for 17 months before scanning.

For MRI, the brain was centered in an acrylic sphere of 6cm diameter (Figure 1A) by gluing the medulla to an alginate socket. The sphere was filled with liquid perfluoropolyether (Fomblin^®^; Solvay Solexis, Bollate, Italy) to protect the specimen from dehydration and to achieve approximate matching of the magnetic susceptibility at tissue interfaces (Benveniste *et al*., 1999).

**Figure 1.**
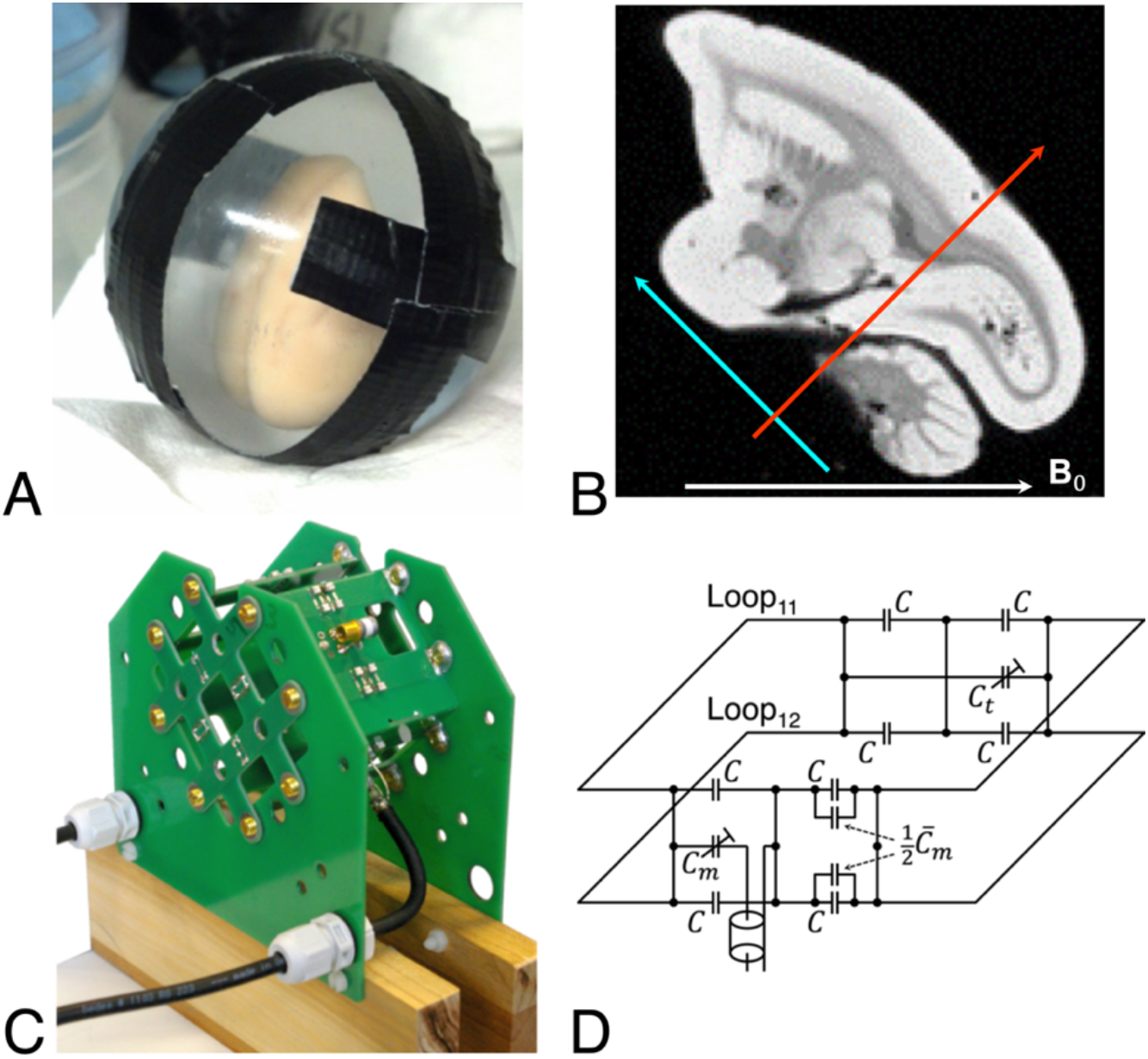
**(A)** Fixed marmoset brain inside a spherical acrylic container filled with Fomblin. **(B)** Sagittal slice from an acquisition at 3 T (flip angle, *α* = 60°; repetition time, *TR* = 300 ms; echo time, *TE* = 20 ms; 200μm isotropic nominal resolution). The direction of the main magnetic field, **B**_0_, is from left to right. Arrows indicate approximate positions and orientations of the horizontal zero plane (blue) and the antero-posterior zero plane (red) in the stereotaxic coordinate system of Paxinos *et al*. (2012). **(C)** Double-Helmholtz transceive RF coil configuration for 3T MRI with additional venting slots and openings for air circulation. The front part is removable for positioning the spherical sample container. Eight non-magnetic brass screws provide electrical contact without degrading the homogeneity of the field amplitude, *B*_0_. Inductive coupling between the perpendicular Helmholtz pairs is negligible for equal currents in both loops of one pair. Holes (8mm diameter) in the PCB at the crossings of the copper traces reduce capacitive coupling caused by mutual capacitances of two pairs (Mispelter *et al*., 2006). **(D)** Tuning (top) and matching (bottom) circuits of a single Helmholtz pair. The feed port is roughly balanced by a matching capacitor (*C*_*m*_) of a few pF. Further improvement is achieved by two capacitors of 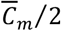 (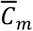 is the average value of *C*_*m*_; *C*_*t*_ is the tuning capacitance).

Using the stereotaxic coordinate system defined by Paxinos *et al*. (2012) as reference (*i.e*., the horizontal, coronal and sagittal zero planes are defined by the plane passing thorough the lower margin of the orbit and the center of the external auditory meatus, the plane passing through the interaural line, and the plane between both hemispheres, respectively), the brain was oriented inside the magnet such that the magnetic field was approximately parallel to the midsagittal plane (*i.e*., azimuthal angle *φ* ≈ 0°) at a polar angle *ϑ* ≈ 45° (Figure 1B).

### 2.2 Magnetic Resonance Image Acquisition

All MRI experiments were performed at room temperature (approx. 21 °C) adjusted by the air-conditioning system of the magnet room, but without an additional temperature control unit for the sample.

#### 2.2.1 Acquisitions at 3 T

A human-scale whole-body scanner (MedSpec 30/100; Bruker Biospin, Ettlingen, Germany) operated under ParaVision 4.0 was used for MRI at 3 T with a custom-built transceiver RF coil. The coil design was made of printed circuit boards (PCBs) and consisted of two perpendicular, quadratic Helmholtz pairs (66×66 mm^2^; 10mm-wide, 35μm-tick copper traces; Figure 1C) to exploit the lower power requirement (*i.e*., reduced sample heating during prolonged scanning) of a quadrature coil (R. Müller *et al*., 2013). Each loop was pre-tuned to 125 MHz by fixed capacitors (*C* = 33 pF; 2%, 1111 SMD footprint, 152 CHB series, Temex Ceramics, Pessac, France), connected in series to ensure balanced feeding (Figure 1D). An additional trimmer capacitor (55H01, Johanson, Boonton, NJ) and the feeding coaxial cable were placed exactly halfway between the loops. A PCB with equivalent layout opposite to the feed port carried the tuning circuit. The Helmholtz pairs were connected to a 90° hybrid via 2m-long coax cables. Lumped resistors of 47 Ω (not related to 50Ω cable impedance) connected the cable sheaths at 150 mm and 400 mm from the coil to achieve broadband damping of parasitic modes (Boskamp *et al*., 2012). Bench-top experiments yielded an unloaded *Q* of 350 and an isolation of the coil pairs by –26 dB. We did not observe relevant detuning, even during scanning sessions of several days.

Experiments in a 50mm-diameter acrylic sphere filled with agarose gel were performed to measure the RF magnetic transmit field, 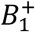. A double-angle method (Hetzer *et al*., 2009; Insko & Bolinger, 1993) with a three-dimensional (3D) Low-Angle SHot (FLASH) sequence (Haase *et al*., 1986) was employed with repetition time TR = 5 s, echo time TE = 6.5 ms, flip angles *α* = 20° and 40°, a field of view (FOV) of 51.2×50×50 mm^3^, and an acquisition matrix 128×50×50. The estimated distribution of 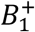 was of sufficient accuracy and homogeneity to allow omitting further corrections of the nominal flip angle (R. Müller *et al*., 2013). Radiofrequency heating experiments (Gaussian pulses; pulse length, *τ*_*p*_ = 10 ms; peak amplitude, *ω*_1,max_ = 18,850 rad/s; TR = 30 ms) performed for 1 hour yielded an increase of the core temperature inside the gel phantom by 6 K as compared to 16 K obtained with a single, linearly polarized Helmholtz coil.

The qMTI protocol was adapted from previous *in-vivo* experiments at 3 T in human subjects (D.K. Müller *et al*., 2013). Magnetization-transfer contrast was generated in a 3D FLASH sequence by applying a preceding 10ms Gaussian ‘MT pulse’ in every repetition (*i.e*., every k-space line). Further acquisition parameters included an ‘imaging pulse’ flip angle, *α* = 10°; TR = 32 ms; and TE = 8.2 ms. To obtain so-called ‘*z*-spectra’ (Grad & Bryant, 1990), a total of 45 image volumes were recorded with different combinations of eleven logarithmically distributed off-resonance frequencies, Ω/(2*π*) = 250–50,000 Hz and seven linearly distributed MT pulse amplitudes with *ω*_1,max_ = 1–7,069 rad/s (Table 1). The FOV was 38.0×27.0×25.6 mm^3^ with a matrix of 190×135×128 (*i.e*., 200μm isotropic nominal resolution). All measurements were averaged 6–16 times, depending on the expected signal-to-noise ratio (SNR) at the particular off-resonance saturation. The total scan time was 72 hours. The first 3 hours of scanning were used to achieve a stable sample temperature. Data acquired during this period were not included in the final analysis. Global *B*_0_ drifts during the experiment were corrected by readjusting the center frequency every 37 to 55 minutes.

**Table 1.** Combinations of MT pulse peak amplitudes, *ω*_1,max_, off-resonance frequencies, Ω/(2π), and numbers of averages, *N*_*a*v_, used for discrete *z*-spectrum sampling. Acquisitions indicated by asterisks were excluded from the final analysis due to reduced accuracy (classifier based on fitting the signal in small water pockets to a single-pool model).

| $\omega_{1,\max}$<br>[rad/s] | $\Omega/(2\pi)$<br>[Hz] | $N_{\text{av}}$ | | $\omega_{1,\max}$<br>[rad/s] | $\Omega/(2\pi)$<br>[Hz] | $N_{\text{av}}$ | |
| --- | --- | --- | --- | --- | --- | --- | --- |
| 1 | 50,000 | 16 |  | 3,770 | 10,353 | 6 |  |
| 1,100 | 250 | 16 |  | 3,770 | 17,624 | 6 |  |
| 1,100 | 426 | 9 |  | 3,770 | 30,000 | 6 |  |
| 1,100 | 724 | 9 |  | 4,712 | 10,353 | 6 |  |
| 1,100 | 1,233 | 9 |  | (5,341 | 250 | 16) | * |
| 1,100 | 2,099 | 6 |  | (5,341 | 426 | 9) | * |
| 1,100 | 3,573 | 6 |  | 5,341 | 724 | 9 |  |
| 1,100 | 6,082 | 6 |  | 5,341 | 1,233 | 9 |  |
| (2,199 | 250 | 16) | * | 5,341 | 2,099 | 6 |  |
| (2,199 | 426 | 9) | * | 5,341 | 3,573 | 6 |  |
| (2,199 | 724 | 9) | * | 5,341 | 6,082 | 6 |  |
| 2,199 | 1,233 | 9 |  | 5,341 | 17,624 | 6 |  |
| 2,199 | 2,099 | 6 |  | 5,341 | 30,000 | 6 |  |
| 2,199 | 3,573 | 6 |  | 7,069 | 426 | 9 |  |
| 2,199 | 6,082 | 6 |  | 7,069 | 724 | 9 |  |
| 2,199 | 10,353 | 6 |  | 7,069 | 1,233 | 9 |  |
| 3,770 | 250 | 16 |  | 7,069 | 2,099 | 6 |  |
| (3,770 | 426 | 9) | * | 7,069 | 3,573 | 6 |  |
| (3,770 | 724 | 9) | * | 7,069 | 6,082 | 6 |  |
| 3,770 | 1,233 | 9 |  | 7,069 | 10,353 | 6 |  |
| 3,770 | 2,099 | 6 |  | 7,069 | 17,624 | 6 |  |
| 3,770 | 3,573 | 6 |  | 7,069 | 30,000 | 6 |  |
| 3,770 | 6,082 | 6 |  |  |  |  |  |

Mapping of the so-called ‘observed’ longitudinal relaxation rate, 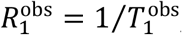, (Henkelman *et al*., 1993) at 3 T was performed with the identical image geometry and nominal resolution as for qMTI with a 3D FLASH sequence (TE = 8 ms) and different combinations of *α* and TR (Fram *et al*., 1987; Helms *et al*., 2008) comprising 10°/30 ms, 20°/30 ms, 30°/30 ms, 30°/90 ms, and 30°/200 ms.

Estimates of the spatial distribution of *B*_0_ across the sample were obtained from two-dimensional (2D) multi-echo (ME) gradient-echo acquisitions (*α* = 60°; TR = 4 s; TE_1_ = 7.79 ms; 32 echoes with inter-echo time ΔTE = 1.28 ms; 32 slices; 800μm nominal isotropic resolution) (Chen & Wyrwicz, 1999; Hetzer *et al*.; 2011).

#### 2.2.2 Acquisitions at 7 T

Further measurements of 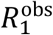 and 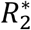 were performed at 7 T on a human-scale whole-body scanner (MAGNETOM 7T; Siemens Healthineers, Erlangen, Germany) operated by *syngo* MR B 17 software. To improve the SNR, a previously described, custom-built miniCP coil was employed, which consisted of two perpendicularly arranged 80-mm circular loops (Weiss *et al*., 2015). Maps of 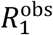 were obtained with the 3D Magnetization-Prepared 2 RApid Gradient Echoes (MP2RAGE) sequence (Marques *et al*., 2010) and parameters (*α*_1_ = *α*_2_ = 8°; TR = 3 s; inversion times, TI_1_ = 250 ms, TI_2_ = 900 ms; TE = 3.43 ms; matrix 160×256×112; nominal resolution 176×176×180 μm; 10 averages) that had been established in former studies of fixed brain tissue (Weiss *et al*., 2015). Maps of 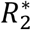 were obtained with a 3D ME-FLASH sequence (*α* = 23°; TR = 42 ms; TE = 6, 14, 22, and 30 ms; matrix 144×192×120; nominal isotropic resolution 200 μm). Finally, 3D FLASH images (*α* = 68°; TR = 0.5 s; TE = 35 ms; FOV 25.88×36×20.8 mm^3^; matrix 506×704×416) were recorded at a high resolution (approx. 50 μm) to improve registration of the magnetic resonance (MR) and histology data by offering sufficiently sharp delineations of tissue boundaries for segmentation and masking purposes.

### 2.3 Magnetic Resonance Image Processing

#### 2.3.1 Image pre-processing

All 3T images were reconstructed offline using in-house software after export of the raw data. Remaining scanner drifts leading to subtle shifts (<1 voxel) of the images along phase-encoding direction were corrected by multiplying appropriate phase ramps to the k-space data (Jenkinson *et al*., 2002; Jenkinson & Smith, 2001).

#### 2.3.2 Magnetization-Transfer Parameter Fitting

In the BSB model, the tissue is subdivided into the free water pool, *‘a’*, and the semisolid pool, *‘b’*, with equilibrium magnetizations 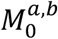 as well as longitudinal and transverse relaxation rates 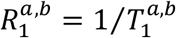 and 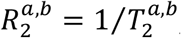, respectively (Edzes & Samulski, 1978; Henkelman *et al*., 1993; Morrison *et al*., 1995). The two pools are further assumed to be in close contact allowing exchange of longitudinal magnetizations 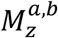 with pseudo-first-order rate constants 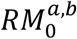 (Henkelman *et al*., 1993). Under these conditions, the time evolution of the magnetization can be described by simplified Bloch-McConnell equations (McConnell, 1958). Finally, saturation of pool *‘b’* caused by the off-resonance irradiation at frequency Ω is modeled by an RF saturation rate (Henkelman *et al*., 1993):

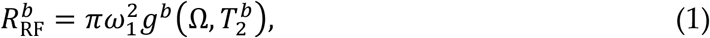

where *ω*_1_ is the RF field amplitude (in rad/s), and 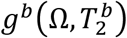 is the absorption lineshape function of the semisolid pool. Consistent with previous work (Morrison *et al*., 1995), we assume that the super-Lorentzian lineshape that arises from partially ordered systems, such as the lipid bilayers of biological membranes (Wennerström, 1973), describes the RF saturation of pool *‘b’* sufficiently well in brain tissue. Finally, a scaling factor, *σ*, is introduced to convert the magnetization computed with the Bloch-McConnell equations into detected signal voltage, 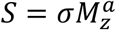.

Exhaustive details of the procedures for BSB parameter fitting have been published elsewhere (D.K. Müller *et al*., 2013). All algorithms were implemented in Matlab 8.1.0.604 (MathWorks, Natick, MA, USA) using the Global Optimization Toolbox (v3.3.1). Unless otherwise stated, least-squares minimization was performed using trust-region-reflective algorithms with parameter boundaries 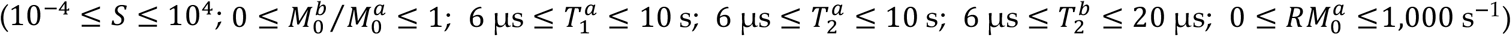. Briefly, fits were based on calculations of the time evolution of the magnetization during the entire pulse sequence using matrix exponentials. The exact timings and shapes of all RF pulses were directly imported from the scanner and used without simplifying assumptions. For better efficiency, polynomial interpolation was employed to calculate the matrix exponentials without bias (D.K. Müller *et al*., 2013; Lenich *et al*., 2019). The separately recorded *B*_0_ map was used for voxel-by-voxel correction of all offset frequencies.

Previous work has shown that oscillations may occur in the *z*-spectrum, particularly at small offset frequencies, which result from the nutation of the liquid-pool magnetization and depend on the MT-pulse amplitude (D.K. Müller *et al*., 2013; Portnoy & Stanisz, 2007). Generally, this effect is difficult to model accurately and may degrade the stability of the fitting procedure. For its further evaluation, the signal amplitude from small pockets of residual water in the alginate socket was fitted to the expected spectrum of a single liquid pool yielding seven data points with residuals outside the 95% confidence interval. Based on this classifier, these seven acquisitions were regarded as potentially affected by oscillations for our experimental conditions and were discarded from the subsequent MT analysis leaving a total of 38 samples in the *z*-spectrum (see Table 1 for details).

In all fits, 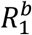 was arbitrarily set to 1 s^−1^ as a fixed parameter. Previous work has shown that its variation over a meaningful range does not lead to an appreciable effect on the *z*-spectrum acquired with steady-state off-resonance irradiation (Henkelman *et al*., 1993; Tyler & Gowland, 2005). As there is a distinct interdependence of some variables, only six BSB model parameters can be uniquely determined from fits to the MT data, namely: 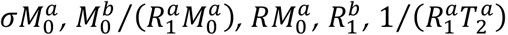, and 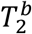 (from 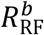).

#### 2.3.3 Estimation of Relaxation Rates

To obtain 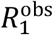 the signal intensities recorded with FLASH and variation of *α* and *TR* were separately fitted to (Ernst *et al*., 1987):

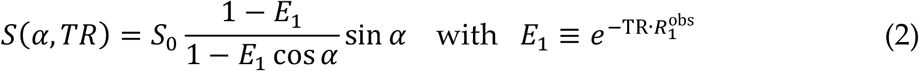

employing a Levenberg-Marquardt algorithm (*S*_0_ is the signal voltage generated by applying a 90° pulse to the fully relaxed spin system). As suggested by Henkelman *et al*., (1993), knowledge of 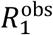 allows computation of 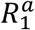 as an additional BSB model parameter.

Estimates of 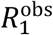 at 7 T were obtained using the vendor software provided with the MP2RAGE sequence (Marques *et al*., 2010) and those of 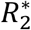 from mono-exponential fits to the TE-dependent signal decay of the ME-FLASH acquisitions.

### 2.4 Histology Procedures

The entire formalin-fixed brain was cut in frozen state into 553 coronal sections of 40μm thickness. During cutting, blockface images (*i.e*., photographs of the top layer of the cutting block) were taken to support volume reconstruction and co-registration of histological and MR data to a mutual reference frame. Every second section was selected for one out of four different staining procedures, which were applied in an alternating fashion (*i.e*., the same staining procedure was applied to every eighth section): *(i)* a modified silver impregnation method to reveal myelin (Gallyas, 1979), *(ii)* immunohistochemical staining for myelin basic protein (anti-MBP, 1:300; Abcam, Cambridge, UK; section immersed in 1% NaBH4 for antigen retrieval), as well as two further antibodies—*(iii)* HuC/HuD antibody (1:500; Life Technologies, Carlsbad, CA, USA) for neurons, and *(iv)* SMI-311 antibody (1:2,000; Calbiochem, San Diego, CA, USA) for pan-neurofilaments—that were not used in the current analysis. In an additional session, several odd-numbered sections were stained for ferric iron using Perls’ stain. Subsequently, the subscripts ‘my’, ‘MBP’ and ‘Fe’ are used to indicate histology results obtained with Gallyas’, MBP and Perls’ stain, respectively.

For a quantitative analysis, initial digitization of multiple slices was performed at relatively low resolution (2.58 μm) on a Zeiss Axio Imager.M1 (Carl Zeiss Microscopy GmbH, Jena, Germany) with an EC Plan-Neofluar 2.5×/0.075 M27 objective and a Zeiss AxioCam HR3 camera. Four sections were then selected (position: approx. 9.75 mm interaural) that showed *(i)* a sufficient variety of anatomical structures and *(ii)* did not indicate major deformations in order to obtain good registration results. Subsequently, maps of the integrated optical density (IOD) were calculated. The sections were digitized monochromatically with 14bit precision, keeping the brightest areas of the imaged window of the sample holder inside the sensitivity range of the CCD sensor. To reduce influences from potential errors due to inhomogeneous dye distribution, the pixel size was set to 0.3225 μm using an EC Plan-Neofluar 20x/0.50 M27 objective (Floyd, 2013). This permits application of the Beer-Lambert law to calculate the IOD or ‘absorbance’:

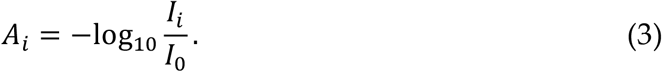

From the mean incident intensity, *I*_0_, measured in an empty area of the sample holder, and the transmitted intensity, *I*_*i*_, at position *i*. The IOD maps were subsequently normalized to a maximum IOD value of 1 in each slice according to:

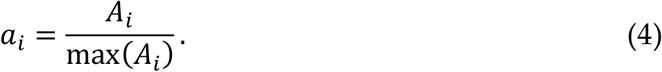

An overview of photographs (at 0.3225 μm) of differently stained slices and corresponding IOD maps is presented in Figure 2.

**Figure 2.**
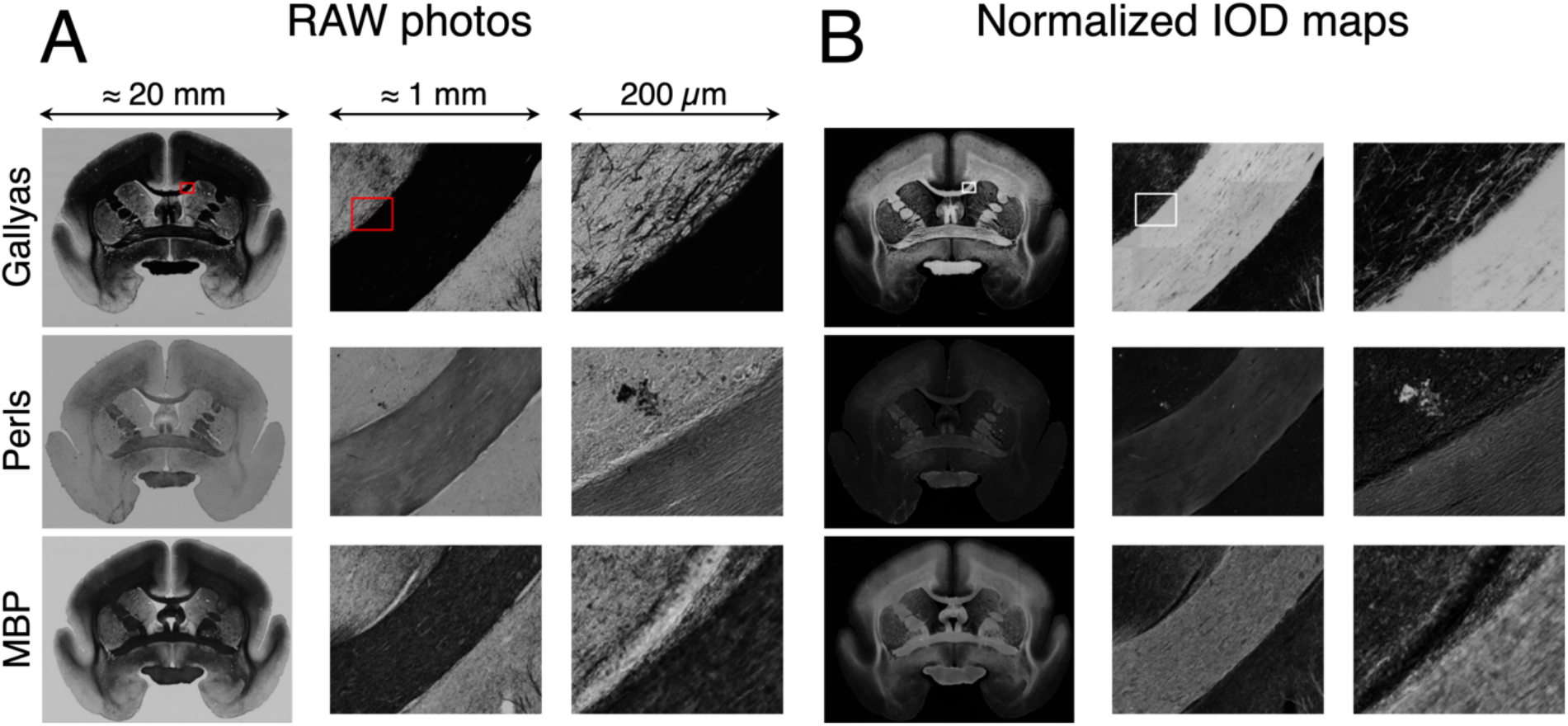
**(A)** Photographs (raw image format) and **(B)** normalized IOD maps (0.3225μm in-plane and 55μm slice resolution) of coronal histology slices at different zoom levels. Rows show, from top (anterior) to bottom (posterior): Gallyas’ silver, Perls’ stain, and anti-MBP immunohistochemistry. An improvement of contrast in WM achieved by calculating normalized IOD maps is evident on slices stained for myelin (Gallyas’ method and anti-MBP immunostaining).

### 2.5 Correlation of MRI and Histology Data

Image registrations were performed using the Image Processing Toolbox (v9.2) of Matlab. The blockface images were concatenated to yield an uncorrected 3D matrix, and the 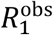 map obtained at 7 T was registered linearly to the blockface reference frame to control for potential misalignment of blockface sections. The low-resolution histological sections were then registered linearly to the appropriate blockface volume sections. The 3T MR parameter maps were registered non-linearly to the 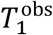 map at 7 T to account for inconsistent geometric distortions due to differences in the individual magnetic field profiles of acquisitions from the different scanners.

As linear registration yielded insufficient accuracy for aligning the high-resolution IOD maps with the MR data, the following non-linear procedure was employed: *(i)* The IOD maps were downsampled (factor of 10) to 3.225 μm and manually segmented into different gray matter (GM) and WM regions. Segmentation was performed by drawing masks along tissue boundaries on highly magnified images using GIMP 2.6.12 (http://www.gimp.org). *(ii)* The high-resolution 7T FLASH images were registered non-linearly to the reference frame while preserving their nominal resolution of 50 μm for sharp tissue boundaries. *(iii)* Slices of the 7T FLASH dataset at positions matching those of the histology slices were segmented into the same regions as done with the IOD maps. *(iv)* Segment maps of the histology data were non-linearly registered to the corresponding segment maps of the 7T FLASH data. *(v)* The resulting deformation fields were used to warp the IOD maps, which were subsequently downsampled to 200 μm. The obtained segment maps were also used in separate analyses of the quantitative MR parameters in GM and WM.

Voxel-by-voxel comparisons of co-registered histology and MR data were limited to a single slice due to concerns of potential variation in the staining intensity between slices (Laule *et al*., 2006). Further quantitative comparisons of different MR parameter maps were performed on the combined data from 21 consecutive coronal sections (region indicated by a red box in the stereotaxic display in Figure 3). This stack of sections also included the single section selected for the comparisons of histology and MR results. To analyze averaged MR parameters in different anatomical structures, segmentations were obtained with FSL 5.0 (Jenkinson *et al*., 2012). The optic chiasm was further segmented from other WM structures for additional investigations.

**Figure 3.**
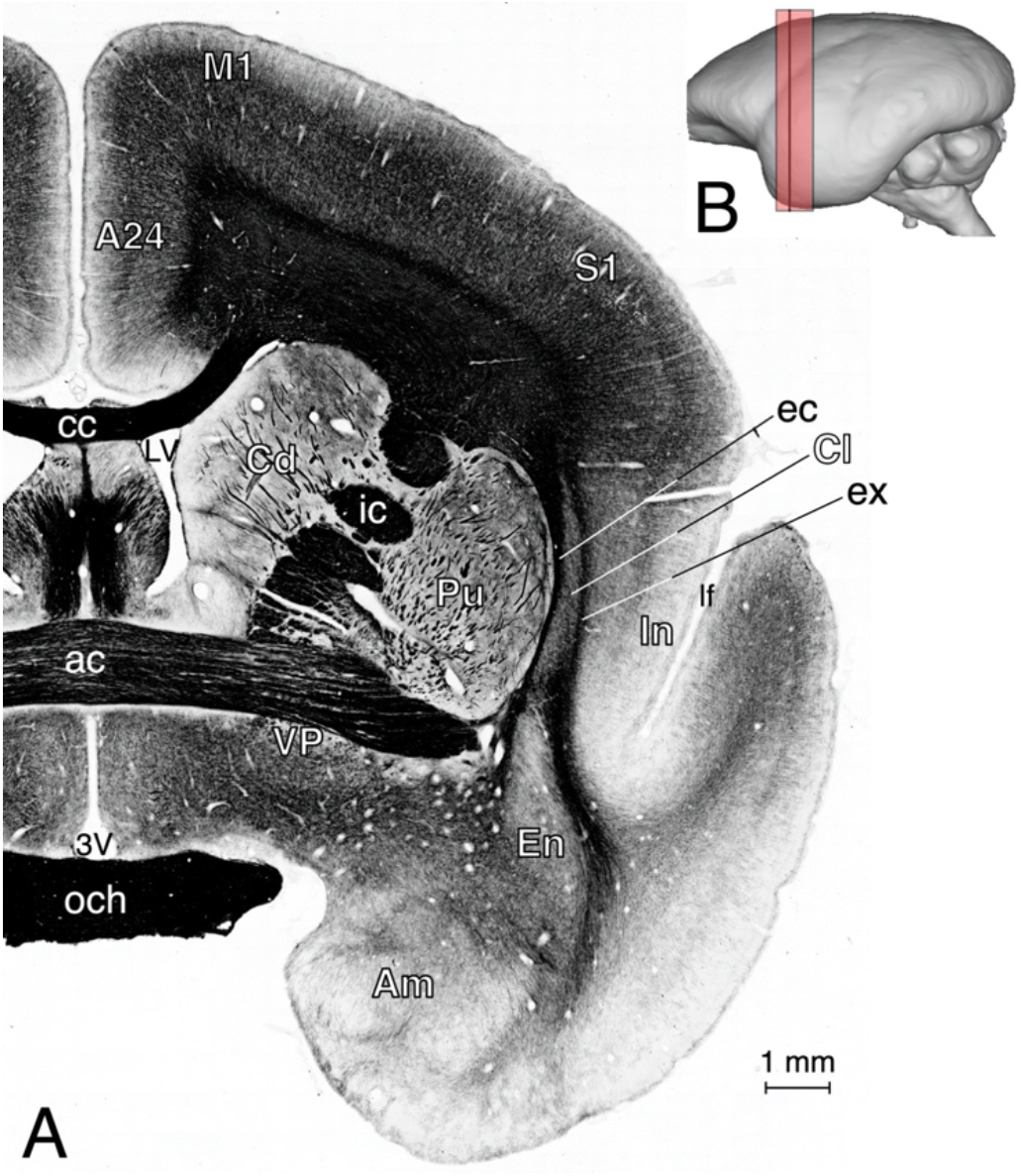
**(A)** High-resolution section of a coronal slice stained for myelin (Gallyas’ silver stain). The slice position is approx. +9.75 mm from the interaural line as marked by a solid line in the stereotaxic view **(B)** [compare, *e.g*., to Figs. 62a and 62 b in (Paxinos *et al*., 2012) or Coronal Plane: Section 19 in (Yuasa *et al*., 2010)]. The red box indicates the position of 21 consecutive coronal slices selected for further voxel-by-voxel correlations of different quantitative MRI parameters. Abbreviations: 3V = 3^rd^ ventricle; A24 = area 24; ac = anterior commissure; Am = amygdaloid nuclei; cc = corpus callosum; Cd = caudate nucleus; Cl = claustrum; ec = external capsule; EN = endopiriform nuclei; ex = extreme capsule; ic = internal capsule; Ins = insula; lf = lateral fissure; LV = lateral ventricle; M1 = primary motor cortex (area 4); och = optic chiasm; Pu = putamen; S1 = primary somatosensory cortex (area 3b); VP = ventral pallidum.

## 3 Results

### 3.1 Histology

All stains yielded stable colorations both across the sections and across the entire volume at visual inspection. For WM areas of highest myelination, the digitization revealed a coloration for the Gallyas silver stain that was close to complete opacity for the procedure employed in the current work. These intensely stained areas showed very low transmitted light intensities approaching the sensor’s electronic noise level, with corresponding IOD values of *A*_my_ < 2. An example is the optic chiasm shown in Figure 3A. To correct for non-zero intensities caused by electronic noise, the minimal brightness of the slice was defined as “pure back”, and all intensity values were shifted accordingly to compute corrected IOD maps.

Voxelwise correlations between the IODs corresponding to different stainings are summarized in Figure 4. Note that these comparisons have an inherently limited accuracy as the different stains were not obtained from identical but from adjacent sections. While this contributes to the scatter in the correlations, in particular in regions of anatomical boundaries, the effect is assumed to be of minor impact as the histology data were downsampled to the much coarser resolution of the MRI acquisitions (200 μm) in these comparisons.

**Figure 4.**
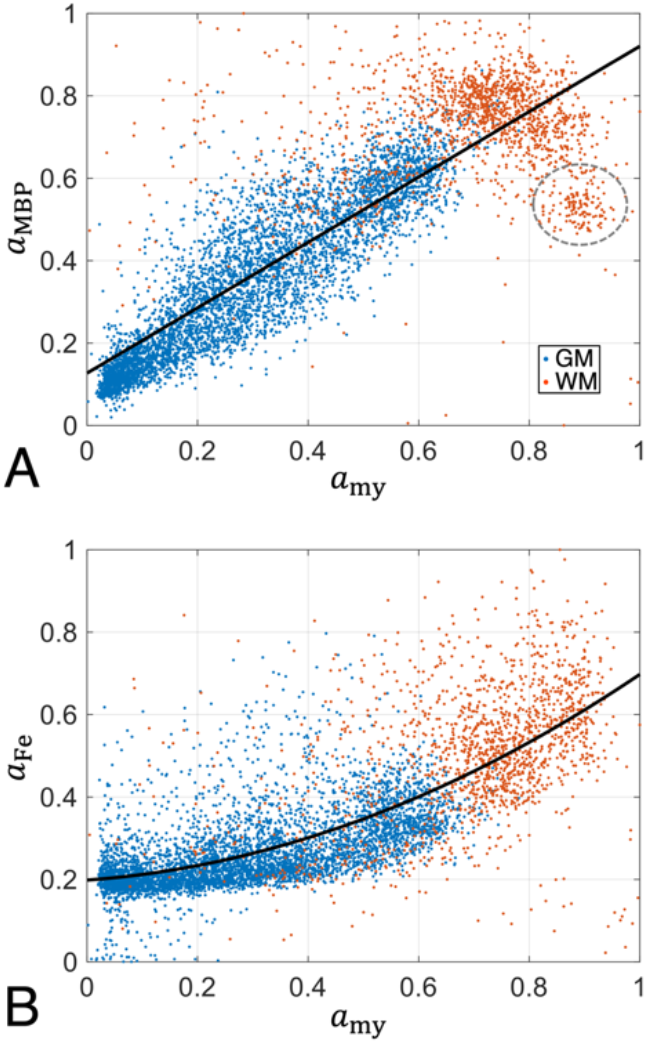
Scatterplots illustrating histology results (normalized IOD) from two myelin stains and Perls’ stain for iron. Blue and red dots indicate voxels in GM (*n* = 4,964) and WM (*n* = 1,538), respectively. Black solid lines show results from polynomial regression analyses. **(A)** A linear correlation of the two myelin stains, *a*_MBP_ = (0.856 ± 0.012) · *a*_my_ + (0.0953 ± 0.0052), was obtained for *a*_my_ < 0.77. Deviations from the regression line were evident for voxels in the highly myelinated optic chiasm (data points enclosed by broken line), which were, therefore, eliminated from the analysis. **(B)** The relation between *a*_Fe_ and *a*_my_ deviated from a linear behavior if GM and WM voxels were included in the same analysis, yielding an approximately quadratic empirical relation with 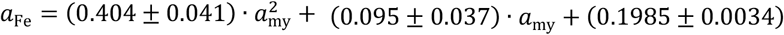.

#### 3.1.1 Comparison of Myelin Stains: Gallyas vs. MBP

A comparison of the normalized IOD values of the two myelin stains revealed a positive linear relation between *a*_my_ and *a*_MBP_ for values covering approximately three quarters of the normalized range, 0 < *a*_my_ < 0.77 (Figure 4A). Anatomical regions with *a*_my_ values in this range included GM in addition to WM structures of medium or lower myelination. In WM structures of highest myelination, such as the optic chiasm, a deviation from the regression line was observed, with *a*_MBP_ values well below those of *a*_my_. This inconsistent behavior became particularly evident when GM and WM regions were analyzed separately, yielding a strong positive correlation in GM (Pearson correlation coefficient, *r* = 0.894; error probability *p* < 0.001, Bonferroni-corrected; see Supplementary Table S1), but an insignificant correlation in WM (*r* = 0.041; *p* = 0.112, uncorrected). Due to the apparently more stable coloration across the entire slice obtained with the silver stain, it was selected for the further analyses.

#### 3.1.2 Comparison of Myelin and Iron Stains

A comparison of normalized IOD values of iron and myelin stains is presented in Figure 4B. A similar behavior was found for *a*_MBP_ (not shown), however, with increased variance compared to *a*_my_, consistent with the results in Figure 4A. Overall, *a*_Fe_ increased with increasing *a*_my_, which could be fitted to an approximately quadratic empirical relation. The observation of a relatively high iron content in WM structures is in line with previous literature demonstrating that oligodendrocytes are the main iron-containing cells in the adult brain (Möller *et al*., 2019; Connor & Menzies, 1996).

### 3.2 Quantitative MRI

No indications of tissue degradation (*e.g*., drifting or inconsistent MR parameters) were observed during the experiments at 3 T and, subsequently, at 7 T. The MR images showed a number of signal voids distributed over the entire volume, which were probably caused by blood clots. The stack of slices used for comparisons of individual MR parameters (Figure 3B) and the single slice used for correlations with histology (Figure 3A) showed a single void located in the left putamen, which was masked out in the quantitative analyses.

Voxelwise fits of the BSB model to the qMT data yielded robust results (Figure 5), with only small variations of the estimated fitting errors across the entire volume. Examples of various MR parameter maps are shown in Figures 6A–C. A good differentiation of GM and WM regions was achieved with the pool-size ratio 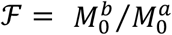, whereas the exchange-rate constant 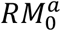 showed a limited dynamic range and, hence, largely uniform intensity across the section. Substantial contrast inside the optic chiasm was observed for the transverse relaxation time of the semisolid pool (Figure 6B), with longer 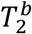 in lateral regions as compared to the central part. The contrast obtained with the relaxation parameters 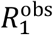 and 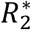 resembled that of ℱ with better SNR (Figures 6D–F). Compared to the 3T result, the 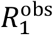 maps obtained at 7T had a higher SNR and improved sharpness of tissue boundaries. Averaged values of MT parameters and results from 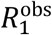 and 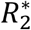 mapping inside selected ROIs in cortical GM, subcortical GM and WM are summarized in Table 2.

**Table 2.** Quantitative results [mean values plus/minus one standard deviation (SD) within the indicated ROI] from MT parameter fitting as well as measurements of the relaxation times 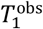 (at 3 T and 7 T) and 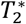. Abbreviations: Am = amygdaloid nuclei; cc: corpus callosum; cr = corona radiata; och, c = optic chiasm, central part; och, l = optic chiasm, lateral part; PaC = parietal cortex; Pu = putamen (see also Figure 3A).

| Region | $\mathcal{F}$ | $RM_0^a$<br>[s<sup>-1</sup>] | $T_1^a$<br>[ms] | $T_2^a$<br>[ms] | $T_2^b$<br>[μs] | $T_1^{\text{obs}}$<br>[ms] | | $T_2^*$<br>[ms] |
| --- | --- | --- | --- | --- | --- | --- | --- | --- |
|  |  |  |  |  |  | 3 T | 7 T |  |
| PaC | 0.13±0.01 | 28.8±2.1 | 347± 9 | 24.7±0.8 | 9.35±0.17 | 376± 8 | 587±15 | 46.8±3.8 |
| Am | 0.10±0.00 | 24.9±1.2 | 423±11 | 30.9±1.3 | 8.45±0.17 | 448±11 | 832±25 | 64.7±9.7 |
| Pu | 0.11±0.00 | 27.3±1.4 | 372±11 | 27.8±0.9 | 8.37±0.17 | 392± 9 | 637±21 | 43.7±7.4 |
| cc | 0.23±0.02 | 27.2±3.4 | 280±33 | 17.7±2.2 | 9.82±0.38 | 319±12 | 439±22 | 25.2±3.2 |
| cr | 0.28±0.01 | 28.2±1.4 | 246±10 | 14.0±0.5 | 10.44±0.25 | 302± 8 | 440±15 | 22.8±1.5 |
| och, c | 0.45±0.04 | 25.9±1.7 | 201±14 | 11.2±0.7 | 9.27±0.32 | 286± 5 | 395±16 | 17.9±2.0 |
| och, l | 0.43±0.04 | 25.6±1.7 | 207±15 | 11.4±1.3 | 10.77±0.32 | 280± 6 | 395±17 | 21.7±1.0 |

**Figure 5.**
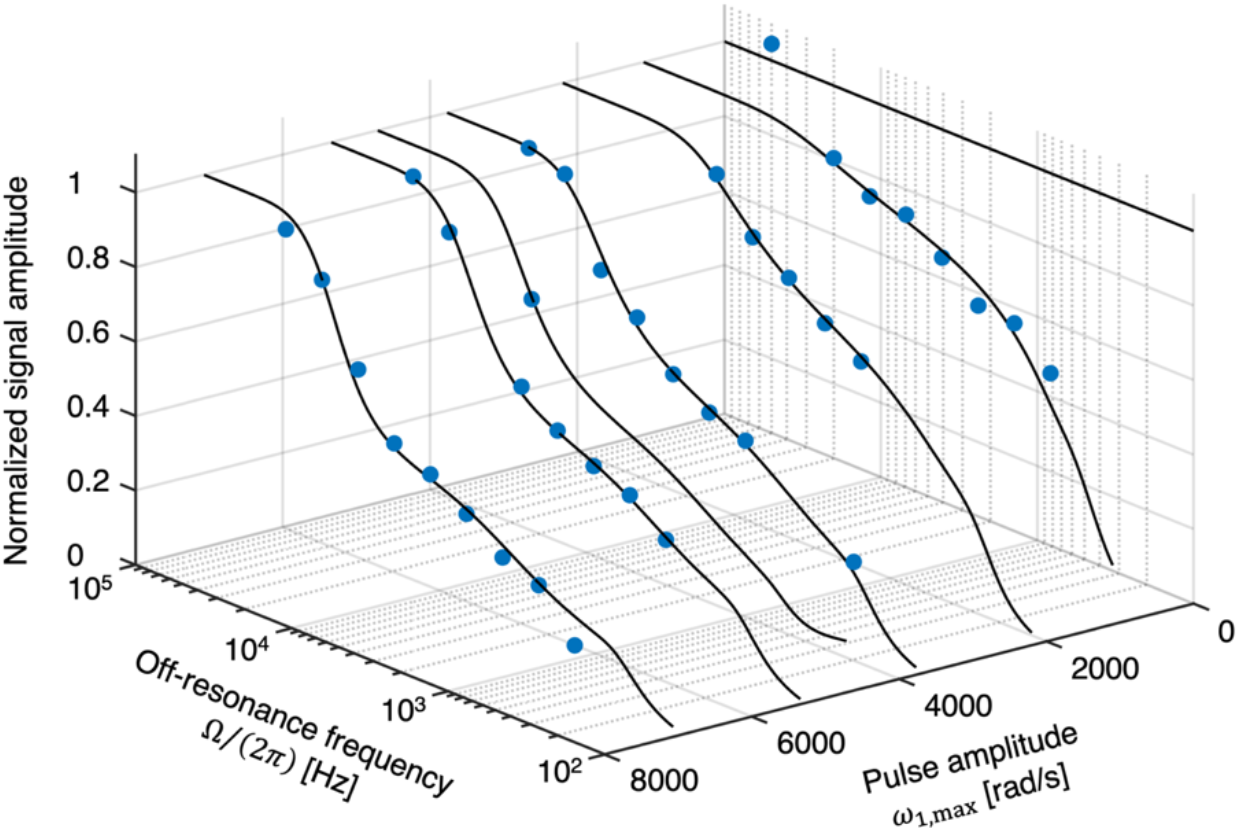
Experimental *z*-spectrum from a single WM voxel consisting of 37 samples remaining after quality assurance (filled blue circles). The steady-state data were acquired with variation of the MT pulse peak amplitude, *ω*_1,max_, and off-resonance frequency, Ω/(2*π*) (see Table 1). Solid black lines show fitting results based on the BSB model. The data are normalized to the maximum signal intensity obtained by the fit in the displayed range (*i.e*., the estimated intensity at 50 kHz off resonance).

**Figure 6.**
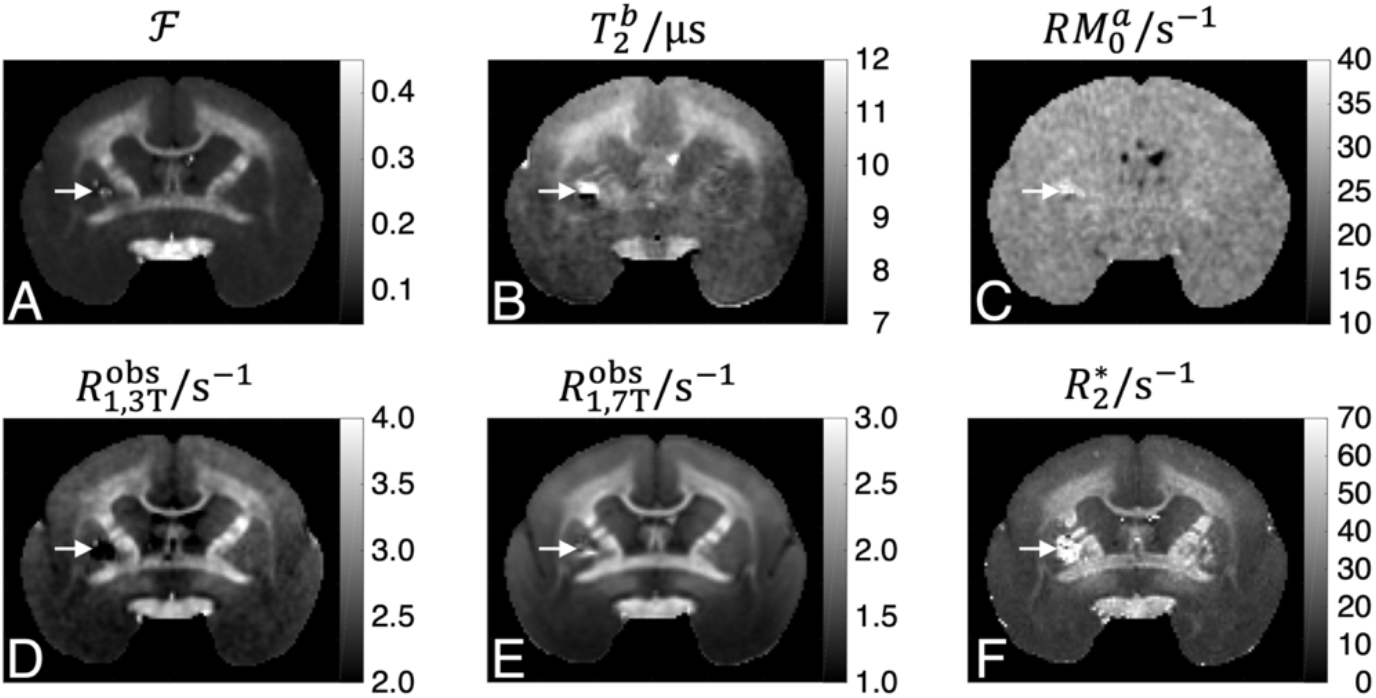
Maps of MT and relaxation parameters including **(A)** the pool-size ratio 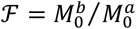, **(B)** the transverse relaxation time of the semisolid pool 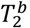, and **(C)** the pseudo-first-order rate contract 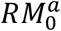, as well as the observed longitudinal relaxation rate 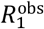 measured at **(D)** 3 T and **(E)** 7 T, and **(F)** the effective transverse relaxation rate 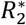 at 7 T. White arrows indicate the position of a signal void in the left putamen, probably due to a blood clot. It is surrounded by a hyperintense area on the 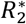 map because the associated field perturbation reaches out into the adjacent tissue. This region was masked out for the further analysis.

A voxel-by-voxel comparison of 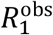 values at the two field strengths yielded a linear relation for the entire value range (*i.e*., including all GM and WM voxels), with rates measured at 3 T exceeding those at 7 T by 30–85% (Figure 7A and Table 2). Plotting 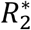 (Figure 7B) as a function of 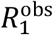 demonstrated deviations from a common regression line when including all tissue classes. Reasonable linear relations, 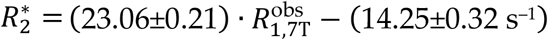 in GM and 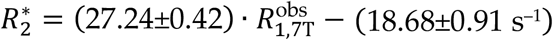 in WM, were obtained in separate analyses of the two tissue classes (Figure not shown).

**Figure 7.**
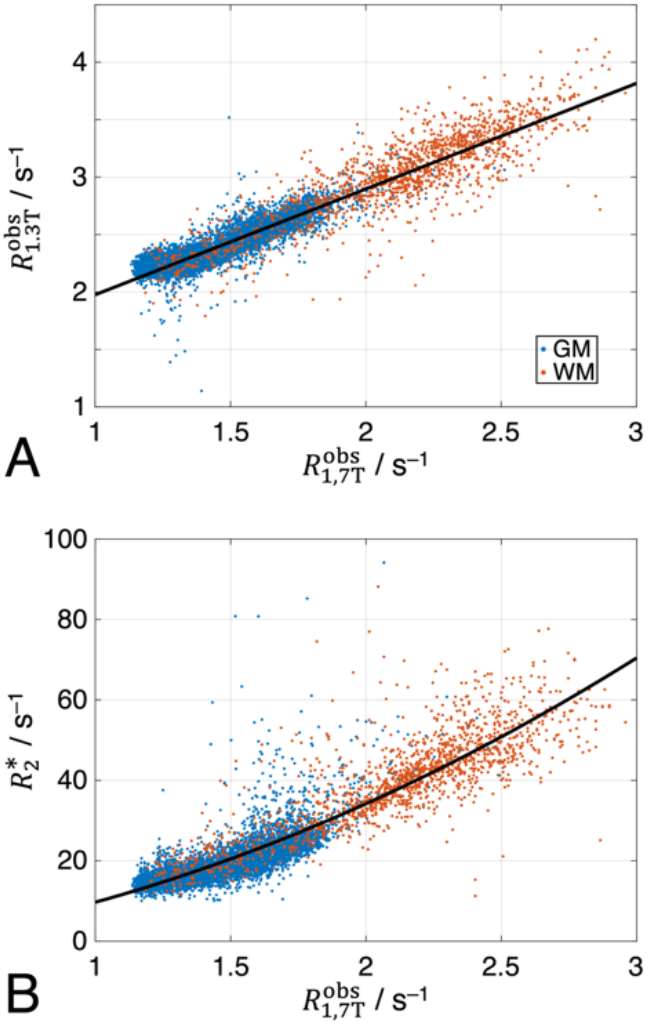
Scatterplots illustrating voxelwise comparisons of the myelin-sensitive MR parameters 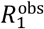 at 3T and 7T and 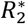 at 7T. Blue and red dots indicate voxels in GM (*n* = 4,964) and WM (*n* = 921), respectively (same locations as in Figure 4). Black solid lines show results from regression analyses: **(A)** 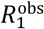 at the two field strengths were linearly correlated for the combined data from GM and WM, 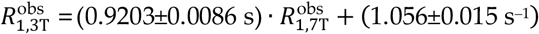. **(B)** The relation between 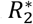 and 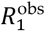 (both measured at 7 T) could be fitted to an approximately quadratic empirical relation, 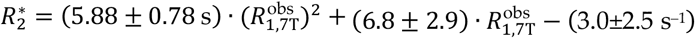.

### 3.3 Correlation of MRI and Tissue Composition Data

The non-linear registration of the multi-modal images enabled a number of explorative univariate analyses on a voxel level, which are summarized in Table 3 and Supplementary Table S1. Strong positive correlations (*p* ≪ 0.001, corrected) with *a*_my_ were obtained for all proposed MR-derived myelin biomarkers 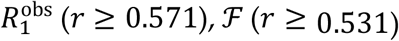 and 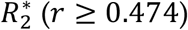 in both GM and WM. Correlations of 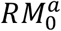 were weaker (*r* ≥ 0.156) but still highly significant. The correlations with *a*_MBP_ were similar to those with *a*_my_ in GM but partly negative or insignificant in WM, which was likely due to inconsistent staining results in regions of highest myelination as mentioned above (see Figure 4A). Furthermore, all MR parameters correlated with *a*_Fe_ yielding a similar range of Pearson coefficients as obtained with *a*_my_.

**Table 3.** Parameter estimates (±SD) from voxelwise univariate linear regressions according to *y*(*x*) =*c*_1_*x* + *c*_0_. Analyses were performed for the whole slice (*n* = 5,885 voxels) shown in Figure 3 and, separately, for masks including only GM (*n* = 4,964 voxels) or only WM (*n* = 921 voxels). Voxels in the optic chiasm were excluded.

| $y(x)$ | $x$ | Whole slice | | GM mask | | WM mask | |
| --- | --- | --- | --- | --- | --- | --- | --- |
| | | $c_1$ | $c_0$ | $c_1$ | $c_0$ | $c_1$ | $c_0$ |
| $a_{\text{MBP}}$ | $a_{\text{my}}$ | 0.8555±0.0058 | 0.1052±0.0025 | 0.8906±0.0045 | 0.0887±0.0016 | -0.082±0.019 | 0.816±0.014 |
| $\mathcal{F}$ | $a_{\text{my}}$ | 0.1493±0.0018 | 0.08368±0.00078 | 0.05904±0.00095 | 0.1033±0.00033 | 0.1508±0.0068 | 0.1213±0.0050 |
| $\mathcal{F}$ | $a_{\text{Fe}}$ | 0.2745±0.0030 | 0.05468±0.00099 | 0.1327±0.0018 | 0.08546±0.00049 | 0.1701±0.0067 | 0.1446±0.0035 |
| $RM_0^a/s^{-1}$ | $\mathcal{F}$ | 16.04±0.59 | 24.68±0.086 | 27.2±1.0 | 23.39±0.13 | 19.2±1.1 | 23.67±0.26 |
| $RM_0^a/s^{-1}$ | $a_{\text{my}}$ | 4.43±0.11 | 25.29±0.049 | 5.41±0.11 | 25.07±0.037 | 4.14±0.38 | 25.08±0.28 |
| $RM_0^a/s^{-1}$ | $a_{\text{Fe}}$ | 6.04±0.21 | 25.06±0.069 | 6.39±0.23 | 24.98±0.064 | 5.58±0.38 | 25.27±0.20 |
| $R_{13T}^{\text{obs}}/s^{-1}$ | $\mathcal{F}$ | 6.021±0.044 | 1.694±0.0064 | 7.161±0.071 | 1.561±0.0086 | 6.16±0.11 | 1.635±0.026 |
| $R_{13T}^{\text{obs}}/s^{-1}$ | $a_{\text{my}}$ | 1.073±0.012 | 2.135±0.0051 | 0.7047±0.0087 | 2.217±0.0030 | 1.349±0.051 | 2.079±0.037 |
| $R_{13T}^{\text{obs}}/s^{-1}$ | $a_{\text{Fe}}$ | 1.957±0.020 | 1.931±0.0065 | 1.346±0.017 | 2.068±0.0049 | 1.773±0.044 | 2.161±0.023 |
| $R_{17T}^{\text{obs}}/s^{-1}$ | $\mathcal{F}$ | 6.095±0.044 | 0.7572±0.0064 | 7.124±0.072 | 0.6319±0.0089 | 4.832±0.097 | 1.054±0.023 |
| $R_{17T}^{\text{obs}}/s^{-1}$ | $a_{\text{my}}$ | 1.135±0.011 | 1.186±0.0047 | 0.7596±0.0083 | 1.267±0.0029 | 1.146±0.041 | 1.338±0.030 |
| $R_{17T}^{\text{obs}}/s^{-1}$ | $a_{\text{Fe}}$ | 1.997±0.019 | 0.9924±0.0063 | 1.369±0.017 | 1.121±0.0049 | 1.44±0.037 | 1.441±0.019 |
| $R_2^*/s^{-1}$ | $\mathcal{F}$ | 177.4±1.5 | -1.15±0.23 | 195.4±2.5 | -3.44±0.31 | 131.2±3.8 | 10.11±0.88 |
| $R_2^*/s^{-1}$ | $a_{\text{my}}$ | 30.67±0.40 | 12.18±0.17 | 16.64±0.31 | 15.23±0.11 | 31.5±1.4 | 17.6±1.0 |
| $R_2^*/s^{-1}$ | $a_{\text{Fe}}$ | 61.36±0.59 | 4.72±0.19 | 46.29±0.52 | 7.85±0.15 | 41.9±1.3 | 19.24±0.68 |

Closer inspection of the correlations between the MR parameters 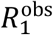 and 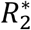 and *a*_my_ indicated characteristic deviations from a straight line for the combined data from GM and WM, whereas the dependencies on *a*_Fe_ were reasonably well described by a common regression line for both segments (Figures 8A,B and 9A,B). Consistent with previous findings (Stüber *et al*., 2014), improved descriptions (*F*-tests; *p* < 0.001, corrected) were obtained with bivariate linear regressions in most cases, according to:

**Figure 8.**
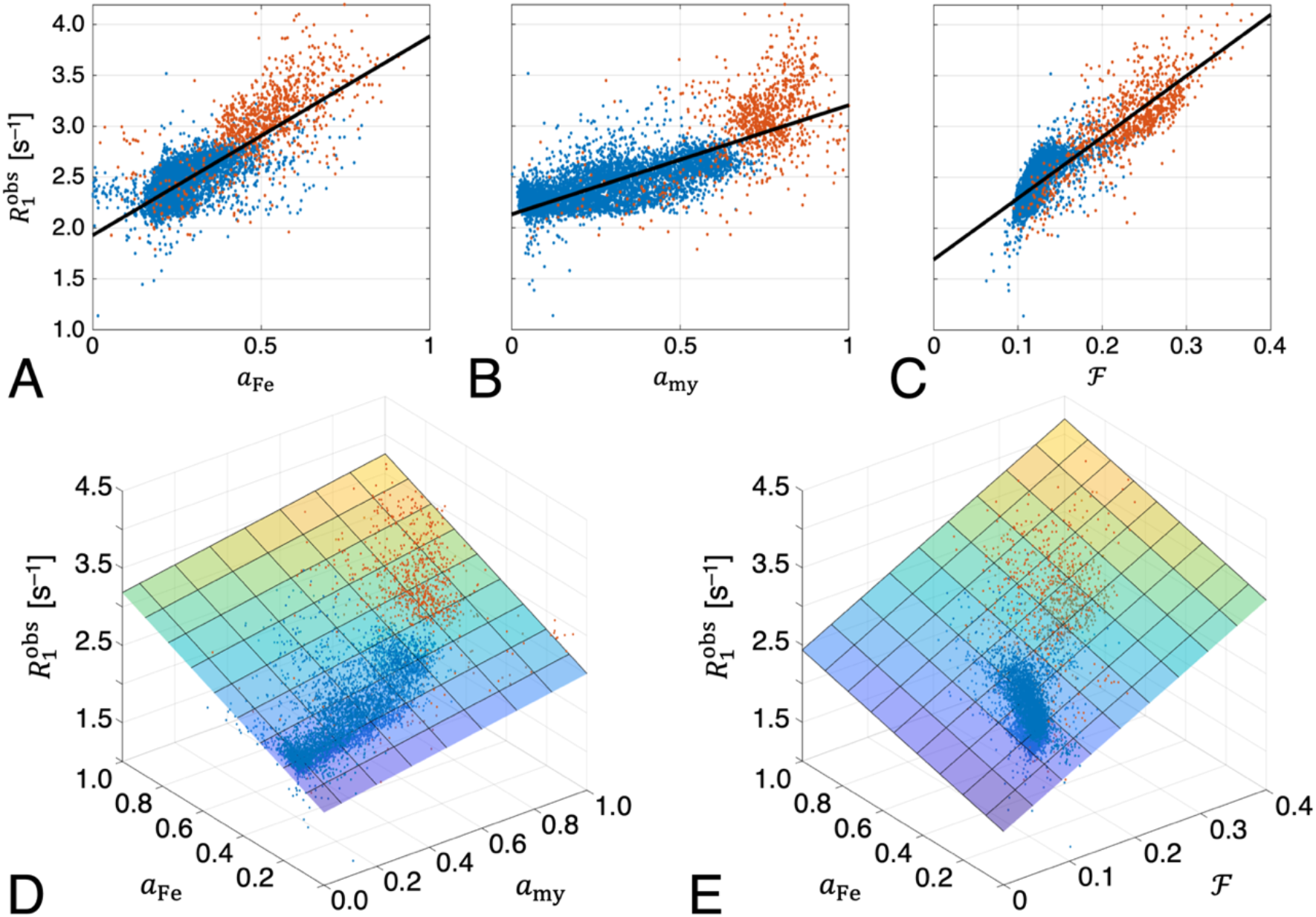
Scatterplots illustrating voxelwise comparisons of the MR relaxometry parameter 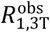 and histology results (normalized IOD) from iron and myelin staining as well as the pool-size ratio ℱ from the MT experiment. Blue and red dots correspond to voxels in GM and WM, respectively. Consistent results were also obtained with 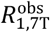. The top row shows results from univariate regressions of **(A)** 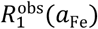, **(B)** 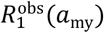 and **(C)** 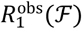. The bottom row shows corresponding bivariate regressions of **(D)** 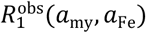 and **(E)** 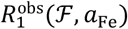 demonstrating significantly improved performance (*p* ≪ 0.001, corrected) compared to the univariate fits 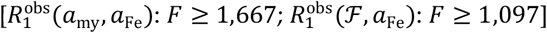.

**Figure 9.**
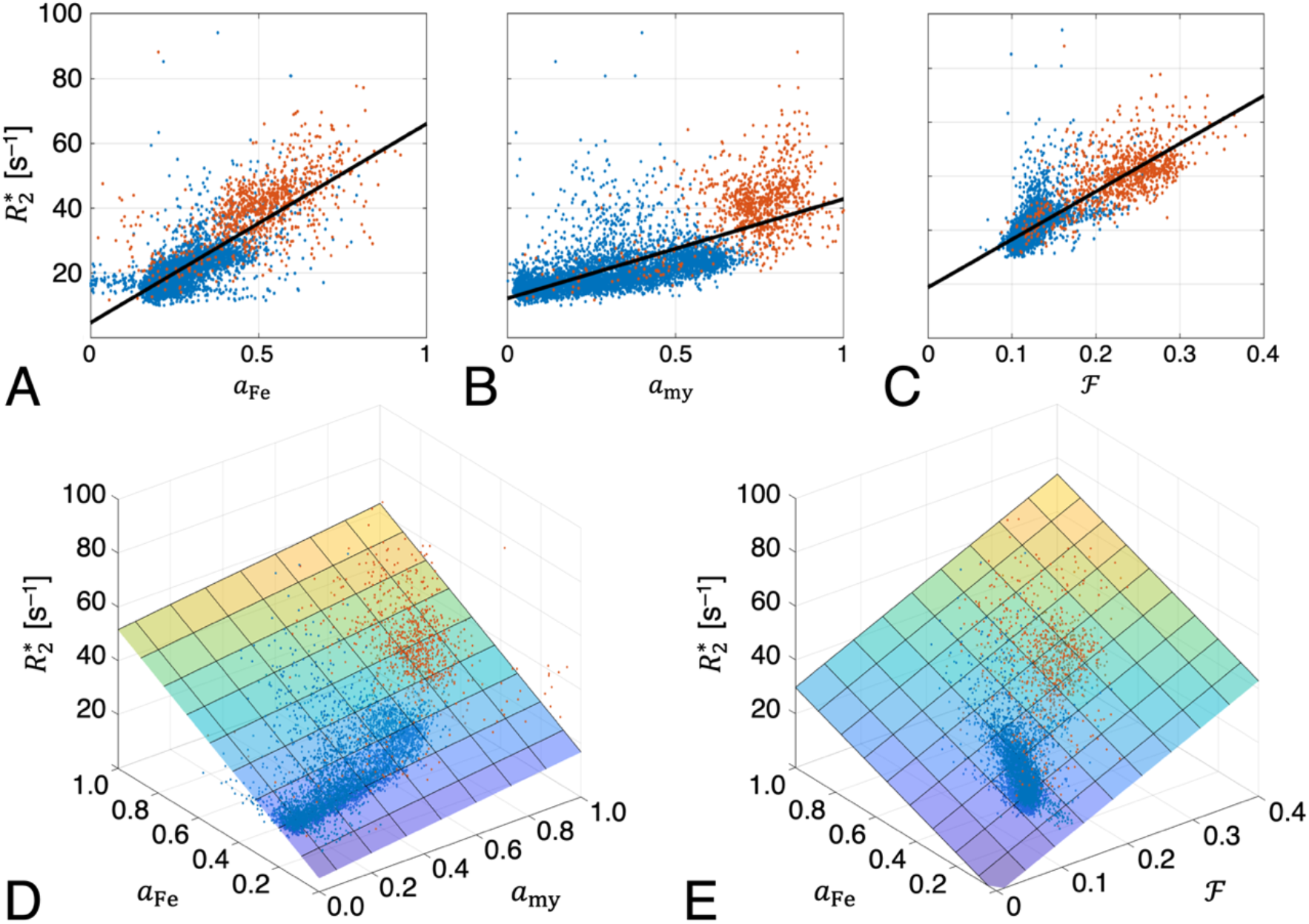
Scatterplots illustrating voxelwise comparisons of the MR relaxometry parameter 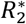 and histology results (normalized IOD) from iron and myelin staining as well as the pool-size ratio ℱ from the MT experiment. Blue and red dots correspond to voxels in GM and WM, respectively. The top row shows results from univariate regressions of **(A)** 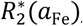, **(B)** 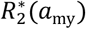 and **(C)** 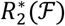. The bottom row shows corresponding bivariate regressions of **(D)** 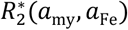 and **(E)** 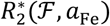 demonstrating significantly improved performance (*p* ≪ 0.001, corrected) compared to the univariate fits 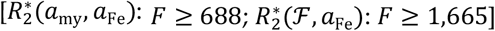.

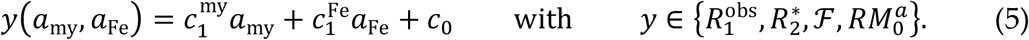

Table 4 summarizes the fitted linear coefficients 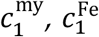 and *c*_0_. Apart from a scaling of the longitudinal relaxation rate reflecting field dependence (see Figure 7A), identical behavior was observed for 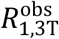 and 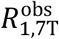.

**Table 4.**
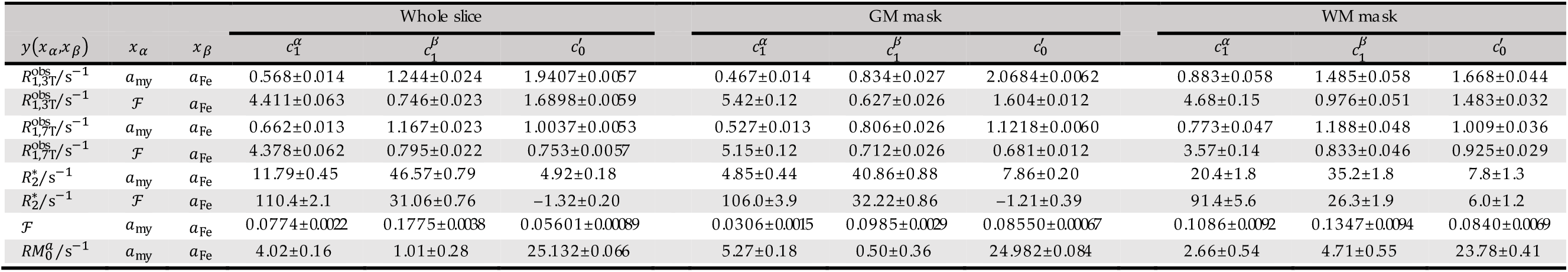
Parameter estimates (±SD) from voxelwise bivariate linear regressions according to 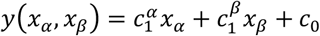. Analyses were performed for the whole slice (*n* = 5,885 voxels) shown in Figure 3 and, separately, for masks including only GM (*n* = 4,964 voxels) or only WM (*n* = 921 voxels). Voxels in the optic chiasm were excluded.

As the simple BSB model does not differentiate between multiple water environments or between multiple macromolecular compartments, a four-pool model was previously proposed for a more comprehensive characterization of brain tissue, and in particular, WM (Barta *et al*., 2015; Bjarnason *et al*., 2005; Levesque & Pike, 2009; Möller *et al*., 2019; Stanisz *et al*. 1999). It suggests that ℱ depends—to first approximation—on contributions from (non-aqueous) myelin and non-myelin dry matter, 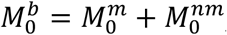, and from myelin water and intra-/extracellular water, 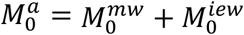. It is further convenient to define corresponding fractions of the total tissue magnetization, 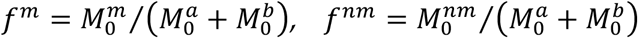 and 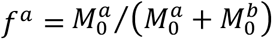, and to express the normalized IOD (Eq. 4) obtained with the Gallyas stain as 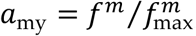, which yields:

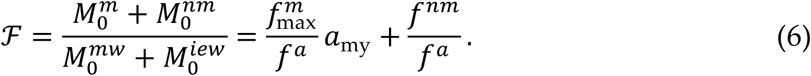

As shown in Appendix A, this leads to an alternative bivariate relation with linear coefficients 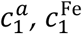 and 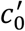

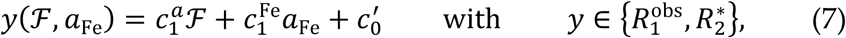

in which the myelin-specific variable *a*_my_ in Eq. 5 is replaced by ℱ combining multiple macromolecular contributions. Note that Eq. 7 was obtained assuming fast intercompartmental water exchange. However, exchange is not sufficiently frequent on the 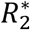 time scale leading to a multiexponential decay (Du *et al*., 2007), which cannot be extracted from our measurement with only four gradient echoes. In case of 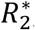. Eq. 7 is, hence, only an empirical relation rather than a model result. Corresponding fits for both relaxation rates are presented in Figures 8C and 9C (results included in Table 4). Compared to results obtained with Eq. 5, the root mean-squared error (RMSE) decreased by 16%, 12% and 12%, and the proportion of the variance explained by the model improved from 71% to 80%, from 76% to 81% and from 69% to 76% for 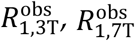 and 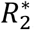, respectively (Supplementary Table S2). This confirms better performance achieved with ℱ comprising multiple water proton relaxants instead of *a*_my_.

A characteristic deviation from a common regression line as observed for the relaxation rates was also evident for the dependency of ℱ on *a*_my_ upon including both GM and WM voxels. Surprisingly, we also obtained a similar dependency of ℱ on *a*_Fe_ (Figure 10A). Finally, 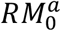 correlated linearly with both *a*_my_ and *a*_Fe_, though without relevant deviations (within the experimental scatter) from common regression lines for the combined GM and WM data (Figures 10D,E). Similar to the relaxation rates, improved descriptions of ℱ were obtained with bivariate linear regression according to Eq. 5 (Figure 10C), whereas a corresponding improvement was small for 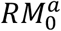 (Figure 10F). In particular, the additional consideration of *a*_Fe_ in 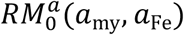 yielded very little improvement compared to a univariate relation 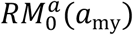 suggesting that any dependence of 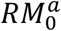 on iron content can only be weak.

**Figure 10.**
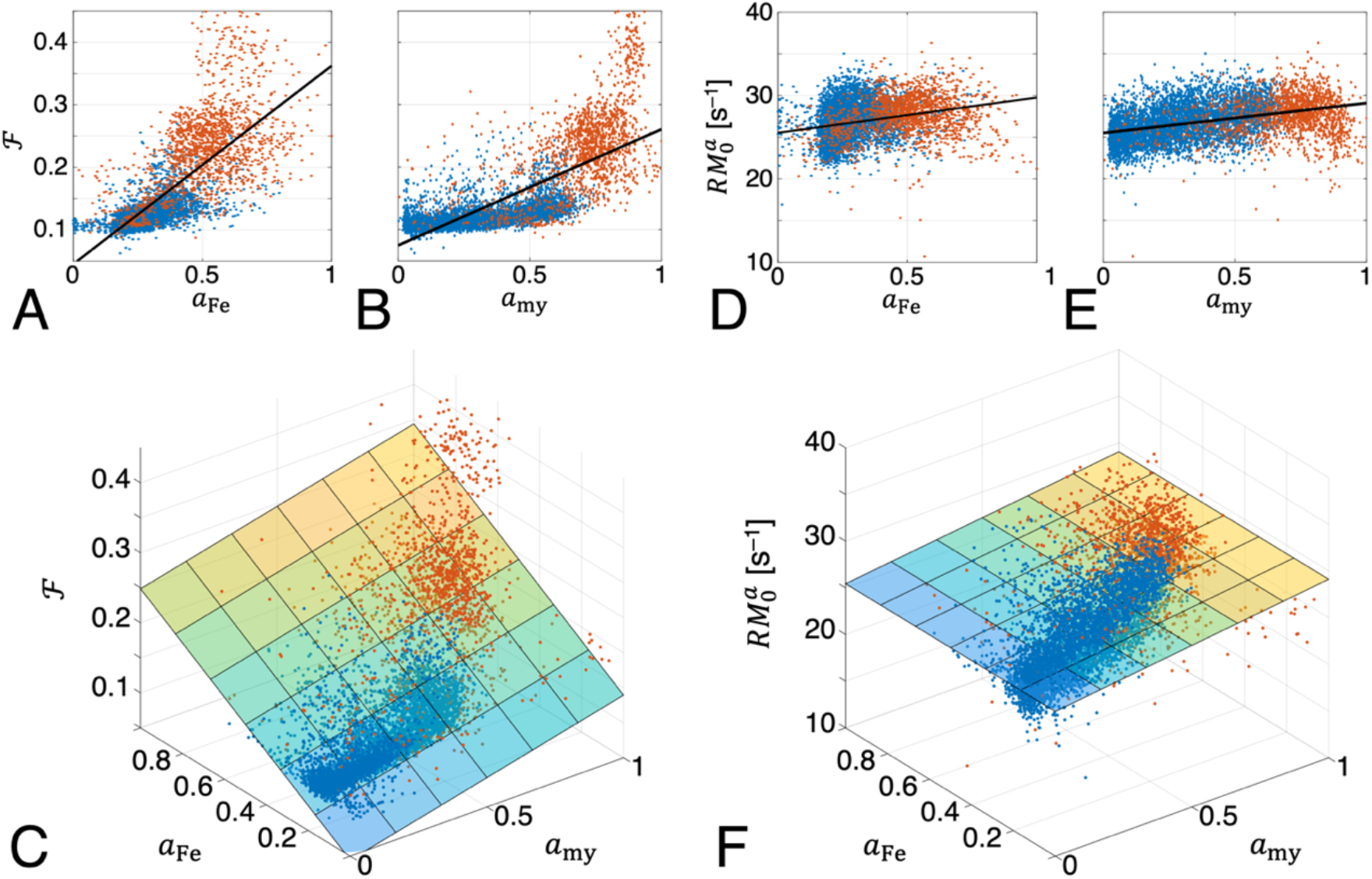
Scatterplots illustrating voxelwise comparisons of the MT parameters ℱ **(A–C)** and 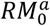 **(D–F)** and histology results (normalized IOD) from iron and myelin stains. Blue and red dots correspond to voxels in GM and WM, respectively. The top row shows results from univariate regressions of **(A)** ℱ(*a*_Fe_) and **(B)** ℱ(*a*_my_) as well as **(D)** 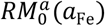 and **(E)** 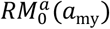. The bottom row shows corresponding bivariate regressions of **(C)** ℱ(*a*_my_, *a*_Fe_) and **(F)** 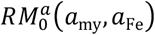. Compared to the univariate fits, performance improved significantly (*p* ≪ 0.001, corrected) for ℱ(*a*_my_, *a*_Fe_) [*F* = 2,137 and 1,255 compared to ℱ(*a*_my_) and ℱ(*a*_Fe_), respectively]. The improvement for 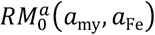 compared to 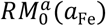 was less pronounced (*F* = 627, *p* ≪ 0.001, corrected) and only marginal (*F* = 13, *p* = 0.006, corrected) when compared to 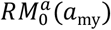.

Comparing the variations of the proposed myelin biomarkers ℱ and 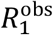 with the histology-derived measure *a*_my_ in GM, a larger (relative) dynamic range is obtained with 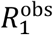 as evidenced by profiles through the primary visual cortex and adjacent WM (Figure 11). The largest intracortical variability was obtained with, 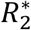 (Figure 11D). This is likely due to the greater sensitivity to the presence of iron for 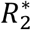 and 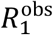 and a presence of iron and myelin in the same voxel (Callaghan *et al*., 2015; Draganski *et al*., 2011; Duyn *et al*., 2007; Helms *et al*., 2008).

**Figure 11.**
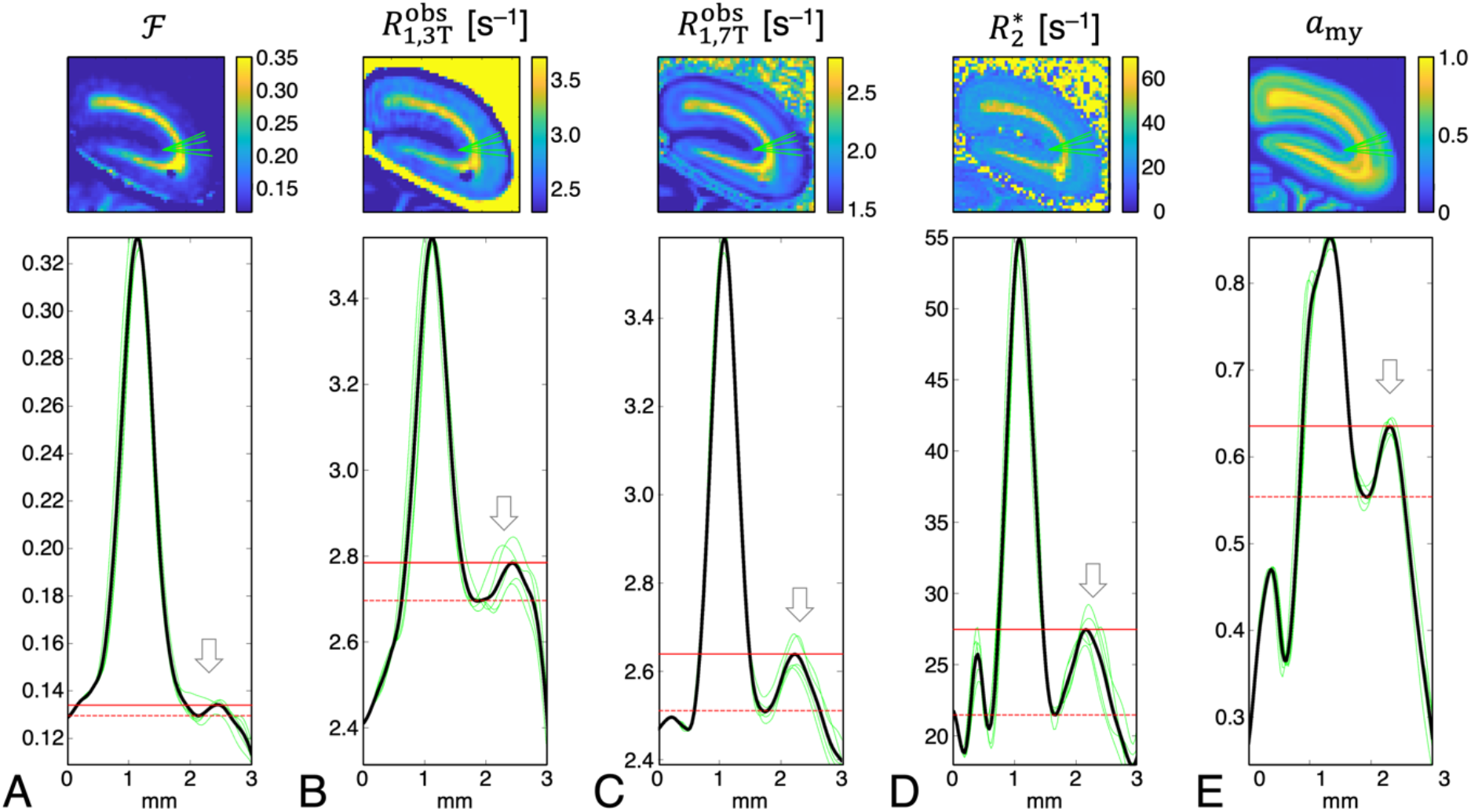
Parameter maps (top row) from a ROI covering parts of the primary visual cortex (V1) and superpositions of profiles (bottom row; light green lines) of **(A)** ℱ, **(B)** 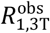, **(C)** 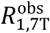, **(D)** 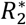 and **(E)** *a*_my_ through V1 and adjacent WM. Solid black lines represent averages of the individual profiles. Positions corresponding to the profiles are indicated by green lines in the parameter maps. The location of the stria of Gennari on the myelin map is indicated by an arrow. The range of variation of the individual parameters between the local maximum inside the stria of Gennari and the local minimum at the GM/WM border is indicated by solid and broken red lines, respectively. The relative parameter variation along the cortical depth is largest for 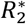, followed by 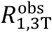 and 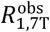 whereas the smallest range is obtained with ℱ.

## 4 Discussion

### 4.1 Comparison of the two Myelin Stains

The mechanism of silver staining of myelin in formalin-fixed brain is assumed to involve reactive foci that bind and reduce silver ions to form metallic clusters, which are visible at microscopy (Larsen *et al*., 2003; Uchihara, 2007). Such foci are ubiquitously present in the myelin sheath. Similarly, MBP is evenly distributed throughout compact myelin (Brunner *et al*., 1989). Hence, both histology methods should report on myelin content. Nevertheless, discrepancies in myelin staining patterns obtained with Gallyas’ method and with anti-MBP immunostaining (or other techniques) have also been observed previously. This includes both GM [*e.g*., (Horton & Hocking, 1997)] and WM [*e.g*., (Kozlowski *et al*., 2008)] as well as a reduced power of resolving fibers in myelin-dense areas with MBP staining in comparison to Gallyas’ method (Pistorio *et al*., 2006). We note that MBP is located in the myelin main period and that low-molecular mass dyes (*e.g*., antibodies to the MBP antigen) do not penetrate the membranes in compact myelin but have to diffuse circumferentially to reach the central laminae of the sheath (Georgi *et al*., 2019; Labadie *et al*., 2014). An apparently reduced IOD observed with anti-MBP immunostaining in WM regions of known high myelination might, therefore, reflect diffusion-limited access of the antibodies with less efficient staining of central laminae during the staining procedure due to later arrival times.

### 4.2 MR Parameters as Biomarkers of Macromolecules and Iron

Our results obtained from a large brain section replicate earlier findings in small tissue specimens, namely bivariate relations between the relaxation rates 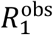 and 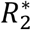 and measures of myelin and iron content (Stüber *et al*., 2014). However, only the dependencies on *a*_Fe_ could be approximated by a common regression line for GM and WM, whereas differences between both tissue segments were evident for the dependencies on *a*_my_. This suggests *(i)* that *a*_Fe_ captures contributions from paramagnetic relaxation enhancement reasonably well and *(ii)* that a restriction to only myelin and iron does not achieve a sufficient explanation of the relaxation effects, hinting at contributions from additional relaxants. Improved fits obtained after replacing *a*_my_ by ℱ corroborate the hypothesis that 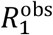 is not uniquely specific to myelination (and iron) but also impacted by other macromolecular factors, such as the density of cells and neuropil. As an additional non-myelin dry-matter fraction was particularly considered in the pool-size ratio in this analysis (Eq. 6), it further suggests that ℱ obtained by qMTI is also not uniquely specific to myelination. In line with this notion, we estimated values of ℱ ≥ 0.1 (Figure 10B) even for very weakly myelinated areas, such as cortical layer II (Niewenhuys, 2012, Tomassy *et al*., 2014; Palomero-Gallagher & Zilles, 2019), confirming that MT is not limited to myelin but includes further contributions.

A ‘nonfreezing’ water component in biological tissues, characterized by slow diffusion with decreased activation energy, is typically identified with a phase of ‘bound water’ (i.e., hydration layers) in models of proton cross-relaxation (Bottomley *et al*., 1984; Escanyé *et al*., 1984; Fullerton *et al*., 1982; Koenig *et al*., 1991). Recent diffusion experiments suggest a substantial contribution from the myelin water fraction (MWF) to the bound pool (Dhital *et al*., 2016). However, myelin water alone failed to account for all of the slowly diffusing component, leaving a portion of 31% in WM that was assigned to water associated with other interfacial structures, which may construct the non-myelin compartment probed in our experiments.

The assumption of a non-myelin macromolecular contribution to ℱ (and 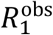) is also in line with earlier observations of different GM/WM ratios derived from different myelin-sensitive MRI techniques: Estimates of the MWF in human GM from multi-exponential *T*_2_-decays were approximately 20% of the values in WM (Laule *et al*., 2007), and 34% in multi-exponential *T*_1_ analyses (Labadie *et al*., 2014). Note that exchange effects cannot be neglected in longitudinal relaxation (Barta *et al*., 2015) and probably contribute to a higher MWF estimate in GM from inversion-recovery data. On the *T*_2_ time scale, the assumption of no water exchange between myelin and intra-/extracellular spaces is reasonable although it may not hold for small, weakly myelinated axons. Therefore, we expect an overall good specificity to the myelin compartment for the MWF. For comparison, previous results for ℱ in human GM *in vivo* were 53% of WM (Sled *et al*., 2004) in agreement with our result of 51% in fixed tissue (parietal cortex vs. corpus callosum and corona radiata, Table 2). This exceeds typical ratios obtained with the MWF by more than a factor of two. The customary MT ratio, defined as MTR = 1 − *S*/*S*_0_, where *S* and *S*_0_ are signal amplitudes measured, respectively, with and without application of an MT saturation pulse, even yielded 69% in GM compared to WM in previous work (Vavasour *et al*., 1998). Besides the ‘true’ MT contribution, which is extracted by a model fit to compute ℱ, the MTR includes another contribution from direct saturation of the water resonance (Henkelman *et al*., 2001). This direct effect increases for longer *T*_1_, amplifying the MTR of GM relative to WM. Consequently, the MTR may be a useful qualitative contrast parameter but is rather limited for extracting quantitative information on macromolecular content.

Further consistency checks are obtained from a closer inspection of the fitting results for 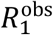 and ℱ:

i. The ratios of the GM and WM slopes and intercepts from univariate fits of ℱ(*a*_my_) to Eq. 6 should be:

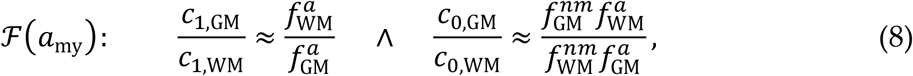

if ℱ includes contributions from myelin and non-myelin dry matter with equal weights. If the non-myelin compartment of WM has a similar composition as that of GM (Norton & Cammer, 1984), we would expect 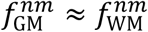. While this is likely an oversimplification, the difference between both fractions is probably small (see Appendix B). The water content is reasonably consistent between mammals and may be estimated from macaque data as 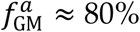 and 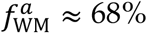 (Faas & Ommaya, 1968; Watanabe *et al*., 1977). This leads to 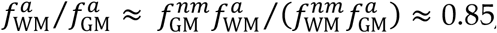, which agrees very well with the observed intercept ratio, *c*_0,GM_/*c*_0,WM_ = 0.852 (Table 3). However, the observed slope ratio, *c*_1,GM_/*c*_1,WM_ = 0.392, deviates from the prediction, which will be further discussed below.
ii. A comparison of Eqs. 5 and 7 (see Appendix A) yields 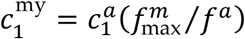, that is, the ratio of the GM and WM slopes from bivariate fits of 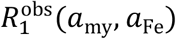 to Eq. 5 should be:

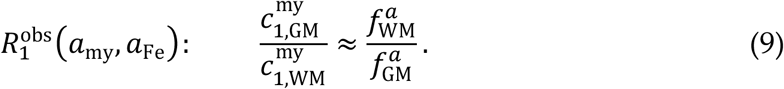

Similar to the result for ℱ(*a*_my_), there is a relevant (though smaller) deviation from the predicted value of 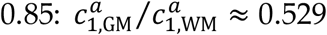 / (Table 4). A potential effect impacting the slopes (Eqs. 8 and 9) could be differences in the lipid composition of GM and WM myelin. Galactolipids, which are more abundant in WM than in GM, have been referred to as most ‘myelin-typical’ lipids (Norton & Cammer, 1984) as their accumulation correlates with the rate of WM myelination during brain maturation (Norton & Poduslo, 1973). They are particularly effective in enhancing water proton relaxation and MT in experiments with model membranes (Kucharczyk *et al*., 1994). Therefore, a potentially higher percentage of galactolipids in WM myelin might amplify the slope in WM. However, this hypothesis is not supported by our results for the exchange rates, 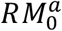, which could be fitted to a common regression line for GM and WM (Fig. 10B). Alternatively, the deviating slopes might suggest different sensitivities of the MT experiment to the myelin and non-myelin compartments. Remarkably, previous multiparametric characterizations of bovine WM based on a four-pool model yielded a more efficient MT exchange rate for the myelin compartment than for the non-myelin compartment (Stanisz *et al*., 1999; Bjarnason *et al*., 2005).
iii. Fits of 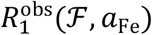 to Eq. 7 should have identical intercepts if further relaxants besides macromolecules and iron are uniformly distributed among tissue classes,

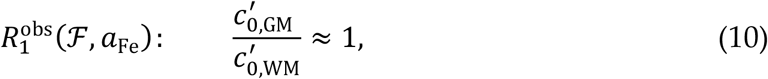

in reasonable agreement with the average experimental results at 3 T and 7 T (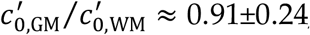; Table 4).

Although previous work has shown that the presence of iron affects the MTR (Smith *et al*., 2009), experiments with the iron storage protein ferritin in model solutions showed that this effect results from water proton *T*_1_ changes rather than an MT effect (Salustri, 1996). Similarly, neuromelanin, that also contains high iron loads, decreased the MTR without an impact on the estimated macromolecular pool size (Trujillo *et al*., 2017). The observed correlations of ℱ and, to a lesser extent, of 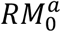 with iron content are, thus, surprising as the pool sizes should not to be directly influenced by iron stores. At the spatial resolution of our MRI experiments (200 μm), myelin and iron are colocalized in the same voxel (Bizzi *et al*., 1990; Duyn *et al*., 2007; Lorio *et al*., 2014; Morris *et al*., 1992), which is reflected in our data in the correlation between *a*_my_ and *a*_Fe_ (Figure 4B). Quantitatively, however, this colocalization did not explain the variations of the MT parameters with *a*_Fe_ well. In this context, a consideration of the different spatial scales of the experiment is important: Both longitudinal relaxation and MT are mediated through dipole-dipole interactions between proton spins and act on a molecular length scale (0.1–1 nm), whereas a cellular scale (1–10 μm) is relevant for the heterogeneous iron distribution (Kiselev & Novikov, 2018). At the cellular level, oligodendrocytes are the most heavily iron-storing cells, followed by microglia, astrocytes and neurons, with approximately three quarters of the total iron being contained in the cytoplasm and mainly localized in lysosomes (Meguro *et al*., 2008; Reinert *et al*., 2019). At a subcellular level, iron accumulation was observed in the inner and outer collars of the myelin sheath but not in compact myelin (Meguro *et al*., 2008). Considering this separation of iron stores and myelin on a micrometer scale, we may assume that histochemical iron measures at the level of an MRI voxel are more associated with membranes of (mostly glial) cell bodies or lysosomes than with myelin. In turn, such a colocalization of iron and a non-myelin macromolecular compartment could produce an apparent correlation of ℱ with *a*_Fe_. We may further speculate that a less efficient MT of non-myelin macromolecules may contribute to the deviations observed for the slope ratios in Eqs. 8 and 9.

### 4.3 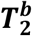 Contrast in the Optic Chiasm

We have recently shown that MT imaging in cerebral WM, and in particular 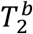, shows an orientation dependence related to the cylindrical symmetry of the myelin membranes enveloping axons (Pampel *et al*., 2015). Previous work has established that the marmoset chiasm is organized as in other primates and humans (Jeffery *et al*., 2008): *(i)* Fibers from the lateral optic nerve (*i.e*., projections from the temporal hemiretina) pass directly (*i.e*., without approaching the midline) through the lateral chiasm toward the ipsilateral optic tract without a change in fiber order. *(ii)* Fibers from the medial optic nerve (*i.e*., projections from the nasal hemiretina) cross the midline in the central chiasm toward the contralateral optic tract. Hence, fibers run in approximately anterior-posterior direction in the lateral chiasm but in approximately left-right (and right-left) direction in the central chiasm. Considering the specimen’s orientation in the magnet, fibers running through the lateral and central parts were approximately at angles of, respectively, 40° and 90° relative to **B**_**0**_. The estimated resulting effect size is consistent with the experimental observation and explains the distinct variation of 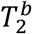 in the optic chiasm. According to previous work (Pampel *et al*., 2015), this orientation effect should not lead to relevant deviations in ℱ and 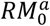.

## Conclusions

Voxel-level comparisons of relaxation rates and MT parameters with histochemical myelin and iron stainings in a whole slice of fixed marmoset brain at high spatial resolution demonstrate high correlations of ℱ, 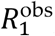, and 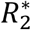 with each other and with the local myelin and iron content. This finding replicates previous results obtained with either larger ROIs or smaller tissue sections of somewhat reduced variability in the compositions. Correlations with the iron content were relatively well described by the same linear dependence for the entire sample, whereas distinct differences were evident in regressions with the myelin content in GM and WM. The combined results suggest that the macromolecular pool impacting relaxation and MT consists of myelin and non-myelin contributions with a more efficient contribution from the myelin compartment. This might be related to the different lipid composition of the two pools, such as, a higher content of galactolipids in myelin. Despite strong correlations of ℱ and 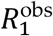 with *a*_my_ content, neither of the two parameters is uniquely specific to myelination because of non-myelin contributions to both MRI-derived biomarkers. Due to the further impact from iron, 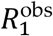 and 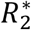 seem more sensitive for depicting microstructural differences between cortical layers. Given that the macromolecular pool is not exclusively from myelin, bias is expected for qMT-derived myelin surrogates, such as MRI-based g-ratio measurements (Stikov *et al*., 2015).

## Appendix A

### Relation between ℱ and 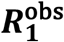

Consistent with previous work, we decompose brain tissue into myelin (*m*) and non-myelin dry matter (*nm*) with proton equilibrium magnetizations 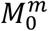 and 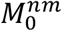, respectively, as well as water within the myelin sheaths (*mw*) and in the remaining intra- and extracellular spaces (*iew*) (Barta *et al*., 2015; Bjarnason *et al*., 2005; Levesque & Pike, 2009; Möller *et al*., 2019; Stanisz *et al*. 1999). Assuming that intra- and extracellular water are indistinguishable by relaxometry, it is sufficient to consider only two water compartments with proton equilibrium magnetizations 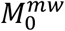 and 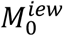, leading to the four-pool model expressed by Eq. 6. Note that ℱ becomes a linear function of *a*_my_ if *f* ^*nm*^ is approximately constant. For this case, the slope depends on *(i)* the maximum myelination observed in the *sample*, 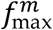, that is, a uniform constant for all GM and WM voxels, and *(ii)* the *voxel’s* water content, *f* ^*a*^, which reflects a *specific tissue type*. In the limit of fast water exchange between the myelin and intra-/extracellular compartments, 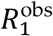 can be written as a linear function of the tissue’s reciprocal water content (Fullerton *et al*., 1982):

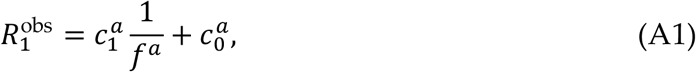

in agreement with experimental results in model systems (Mezer *et al*., 2013) and *in vivo* (Gelman *et al*., 2001). However, Eq. A1 does not consider iron-related relaxation, which may be approximated by a separate linear term. With *f* ^*a*^ = (1 + ℱ) ^− 1^, this leads to the empirical relation (Callaghan *et al*., 2015; Rooney *et al*., 2009):

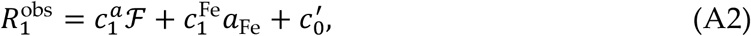

where 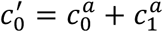. Finally, upon insertion of Eq. 6, we obtain:

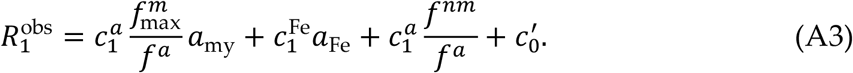

A comparison of Eqs. 5 and A3 yields 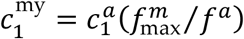 and 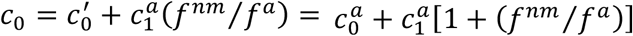. Note that a uniform bivariate relation according to Eq. 5 with identical relaxivities 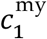 for GM and WM would only result if the water content does not differ between the tissue classes.

## Appendix B

### Lipid Pools in GM and WM

According to Norton and Cammer (1984), fresh human WM contains roughly 300 mg/g dry matter with approximately equal myelin and non-myelin contributions and 700 mg/g water. Of the total water, about 100 mg/g are in the myelin and 600 mg/g in the non-myelin compartment, yielding MWF_WM_ ≈ 0.14. For simplicity, we further assume that cross-relaxation effects, and hence, 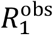 and ℱ, are dominated by (membrane-bound) lipids rather than proteins (Koenig, 1991, Kucharczyk *et al*., 1994; Pampel *et al*., 2015). Of the myelin solids, 30% are proteins and 70% or 105 mg/g are lipids, whereas the lipid fraction of the total dry matter is 54.9% or 165 mg/g. This suggests that myelin accounts for 64% of the lipids and non-myelin for the remaining 36% leading to 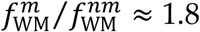. Finally, galactolipids are specifically enriched in myelin (Norton & Poduslo, 1983) and particularly efficient cross-relaxation sites (Kucharczyk *et al*., 1994). They contribute 27.7% of all myelin lipids and 26.4% to the total WM lipid fraction, which corresponds to 29 mg/g and 43 mg/g, respectively. Therefore, 67% of all WM galactolipids are in the myelin and 33% in the non-myelin compartment. Fresh human GM contains about 820 mg/g water and 180 mg/g solids, of which 32.7% or 59 mg/g are lipids. A contribution of 7.3% to the total lipid content is from galactolipids, corresponding to 4.3 mg/g, which defines the maximum possible galactolipid content of GM myelin. An alternative assumption of the same distribution of galactolipids between myelin (67%) and non-myelin (33%) in GM as in WM would lead to approximately 2.9 mg/g. If GM and WM myelin are of similar composition, we expect that 27.5% of all lipids in myelin are galactolipids, and the limiting cases of 2.9– 4.3 mg/g computed above yield 10–15 mg/g myelin water and 15–22 mg/g myelin solids (lipids plus proteins), of which 10–16 mg/g are due to myelin lipids. This leads to MWF_GM_ ≈ 0.012–0.018 and 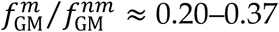. Further comparisons of the estimates for GM and WM yield 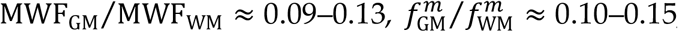, and 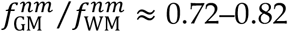. Considering the variation of published values of the lipid composition (Norton & Cammer, 1984; O’Brian & Sampson, 1965), we should consider potential deviations from the above estimates in the order of 10% for the MWF and 25% for the lipid fractions.

## Data and code availability statement

Pre-processed MRI and histology raw data and derived MT and relaxometry parameter maps will be made available upon acceptance at https://dataverse.harvard.edu/. The pre-processing and analyses steps have been well documented in the Methods section. If required, the Matlab scripts used for this purpose can be made available upon request.

## CRediT authorship contribution statement

**Henrik Marschner:** Methodology, Software, Formal analysis, Investigation, Data curation, Writing – review & editing. **André Pampel:** Conceptualization, Methodology, Software, Validation, Investigation, Writing – review & editing. **Roland Müller:** Methodology, Investigation, Writing – review & editing. **Katja Reimann:** Investigation, Data curation, Writing – review & editing. **Nicolas Bock:** Resources, Writing - review & editing. **Markus Morawski:** Conceptualization, Resources, Writing – review & editing. **Stefan Geyer:** Conceptualization, Resources, Writing – review & editing. **Harald E. Möller:** Conceptualization, Methodology, Resources, Writing – original draft, Writing – review & editing, Supervision, Project administration, Funding acquisition.

## Acknowledgements

We thank Afonso Silva for kindly providing the specimen, Marcel Weiss for supporting the 7T acquisitions and Nico Scherf for helpful discussions on analysis strategies. This work was funded (in part) by the Helmholtz Alliance “ICEMED— Imaging and Curing Environmental Metabolic Diseases” (HA-314) through the Initiative and Networking Fund of the Helmholtz Association and by the Max Planck Society.

## References

1. Barta, R., Kalantari, S., Laule, C., Vavasour, I.M., MacKay, A.L., Michal, C.A., 2015. Modeling T_1_ and T_2_ relaxation in bovine white matter. J. Magn. Reson. 259: 56–67. https://doi.org/10.1016/j.jmr.2015.08.001.

2. Benveniste, H., Einstein, G., Kim, K.R., Hulette, C., Johnson, G.A., 1999. Detection of neuritic plaques in Alzheimer’s disease by magnetic resonance microscopy. Proc. Natl. Acad. Sci. USA 96: 14079–14084. https://doi.org/10.1073/pnas.96.24.14079.

3. Bizzi, A., Brooks, R.A., Brunetti, A., Hill, J.M., Alger, J.R., Miletich, R.S., Francavilla, T.L., Di Chiro, G., 1990. Role of iron and ferritin in MR imaging of the brain: A study in primates at different field strengths. Radiology 177: 59–65. https://doi.org/10.1148/radiology.177.1.2399339.

4. Bjarnason, T.A., Vavasour, I.M., Chia, C.L.L., MacKay, A.L., 2005. Characterization of the NMR behavior of white matter in bovine brain. Magn. Reson. Med. 54: 1072–1081. https://doi.org/10.1002/mrm.20680.

5. Bock, N.A., Kocharyan, A., Liu, J.V., Silva, A.C., 2009. Visualizing the entire cortical myelination pattern in marmosets with magnetic resonance imaging. J. Neurosci. Methods 185: 15–22. https://doi.org/10.1016/j.jneumeth.2009.08.022.

6. Boskamp, E.B., Fujimoto, M., Amjad, A., Edwards, M., 2012. Broadband damping of cable modes. In Proceedings of the 20th Annual Meeting of ISMRM, Melbourne, VIC, Australia. p. 2691.

7. Bottomley, P.A., Foster, T.H., Argersinger, R.E., Pfeifer, L.M., 1984. A review of normal tissue hydrogen NMR relaxation times and relaxation mechanisms from 1–100 MHz: Dependence on tissue type, NMR frequency, temperature, species, excision, and age. Med. Phys. 11: 425–448. https://doi.org/10.1118/1.595535.

8. Brunner, C., Lassmann, H., Waehneldt, T.V., Matthieu, J.-M., Linington, C., 1989. Differential ultrastructural localization of myelin basic protein, myelin/oligodendroglial glycoprotein, and 2’,3’-cyclic nucleotide 3’-phosphodiesterase in the CNS of adult rats. J. Neurochem. 52: 296–304. https://doi.org/10.1111/j.1471-4159.1989.tb10930.x.

9. Bryant, R.G., Korb, J.-P., 2005. Nuclear magnetic resonance and spin relaxation in biological systems. Magn. Reson. Imaging 23: 167–173. https://doi.org/10.1016/j.mri.2004.11.026.

10. Callaghan, M.F., Freund, P., Draganski, B., Anderson, E., Cappelletti, M., Chowdhury, R., Diedrichsen, J., FitzGerald, T.H.B., Smittenaar, P., Helms, G., Lutti, A., Weiskopf, N., 2014. Widespread age-related differences in the human brain microstructure revealed by quantitative magnetic resonance imaging. Neurobiol. Aging 35: 1862–1872. https://doi.org/10.1016/j.neurobiolaging.2014.02.008.

11. Callaghan, M.F., Helms, G., Lutti, A., Mohammadi, S., Weiskopf, N., 2015. A general linear relaxometry model of R# using imaging data. Magn. Reson. Med. 73: 1309–1314. https://doi.org/10.1002/mrm.25210.

12. Ceckler, T.L., Wolff, S.D., Yip, V., Simon, S.A., Balaban, R.S., 1992. Dynamic and chemical factors affecting water proton relaxation by macromolecules. J. Magn. Reson. 98: 637–645. https://doi.org/10.1016/0022-2364(92)90018-3.

13. Chen, N.-k., Wyrwicz, A.M., 1999. Correction for EPI distortions using multi-echo gradient-echo imaging. Magn. Reson. Med. 41: 1206–1213. https://doi.org/10.1002/(sici)1522-2594(199906)41:6<1206::aid-mrm17>3.0.co;2-l.

14. Connor, J.R., Menzies, S.L., 1996. Relationship of iron to oligodendrocytes and myelination. Glia 17: 83–93. https://doi.org/10.1002/(SICI)1098-1136(199606)17:2<83::AID-GLIA1>3.0.CO;2-7.

15. Dhital, B., Labadie, C., Stallmach, F., Möller, H.E., Turner, R., 2016. Temperature dependence of water diffusion pools in brain white matter. NeuroImage 127: 135–143. https://doi.org/10.1016/j.neuroimage.2015.11.064.

16. Draganski, B., Ashburner, J., Hutton, C., Kherif, F., Frackowiak, R.S.J., Helms, G., Weiskopf, N., 2011. Regional specificity of MRI contrast parameter changes in normal ageing revealed by voxel-based quantification (VBQ). NeuroImage 55: 1423–1434. https://doi.org/10.1016/j.neuroimage.2011.01.052.

17. Du, Y.P., Chu, R., Hwang, D., Brown, M.S., Kleinschmidt-DeMasters, B.K., Singel, D., Simon, J.H., 2007. Fast multislice mapping of the myelin water fraction using multicompartment analysis of T* decay at 3T: A preliminary postmortem study. Magn. Reson. Med. 58: 865–870. https://doi.org/10.1002/mrm.21409.

18. Duyn, J.H., van Gelderen, P., Li, T.-Q., de Zwart, J.A., Koretsky, A.P., Fukunaga, M., 2007. High-field MRI of brain cortical substructure based on signal phase. Proc. Natl. Acad. Sci. USA 104: 11796–11801. https://doi.org/10.1073/pnas.0610821104.

19. Edzes, H.T., Samulski, E.T., 1977. Cross relaxation and spin diffusion in the proton NMR of hydrated collagen. Nature 265 (5594): 521–523. https://doi.org/10.1038/265521a0.

20. Edzes, H.T., Samulski, E.T., 1978. The measurement of cross-relaxation effects in the proton NMR spin-lattice relaxation of water in biological systems: Hydrated collagen and muscle. J. Magn. Reson. 31: 207–229. https://doi.org/10.1016/0022-2364(78)90185-3.

21. Ernst, R.R., Bodenhausen, G., Wokaun, A., 1987. Principles of Nuclear Magnetic Resonance in One and Two Dimensions. Oxford: Clarendon Press; pp. 124–125.

22. Escanyé, J.M., Canet, D., Robert, J., 1984. Nuclear magnetic relaxation studies of water in frozen biological tissues. Cross-relaxation effects between protein and bound water protons. J. Magn. Reson. 58: 118–131. https://doi.org/10.1016/0022-2364(84)90011-8.

23. Faas, F.H., Ommaya, A.K., 1968. Brain tissue electrolytes and water content in experimental concussion in the monkey. J. Neurosurg. 28: 137–144. https://doi.org/10.3171/jns.1968.28.2.0137.

24. Floyd, A.D., 2013. Quantitative data from microscopic specimens. In: Suvarna, S.K., Layton, C., Bancroft, J.D. (editors). Bancroft’s Theory and Practice of Histological Techniques. 7th Edition. London: Churchill Livingstone; pp. 40–60.

25. Fralix, T.A., Ceckler, T.L., Wolff, S.D., Simon, S.A., Balaban, R.S., 1991. Lipid bilayer and water proton magnetization transfer: Effect of cholesterol. Magn. Reson. Med. 18: 214–223. https://doi.org/10.1002/mrm.1910180122.

26. Fram, E.K., Herfkens, R.J., Johnson, G.A., Glover, G.H., Karis, J.P., Shimakawa, A., Perkins, T.G., Pelc, N.J., 1987. Rapid calculation of T1 using variable flip angle gradient refocused imaging. Magn. Reson. Imaging 5: 201–208. https://doi.org/10.1016/0730-725x(87)90021-x.

27. Fullerton, G.D., Potter, J.L., Dornbluth., N.C., 1982. NMR relaxation of protons in tissues and other macromolecular water solutions. Magn. Reson. Imaging 1: 209–228. https://doi.org/10.1016/0730-725x(82)90172-2.

28. Gallyas, F., 1979. Silver staining of myelin by means of physical development. Neurol. Res. 1: 203–209. https://doi.org/10.1080/01616412.1979.11739553.

29. Gelman, N., Ewing, J.R., Gorell, J.M., Spickler, E.M., Solomon, E.G., 2001. Interregional variation of longitudinal relaxation rates in human brain at 3.0 T: Relation to estimated iron and water contents. Magn. Reson. Med. 45: 71–79. https://doi.org/10.1002/1522-2594(200101)45:1<71::aid-mrm1011>3.0.co;2-2.

30. Georgi, J., Metere, R., Jäger, C., Morawski, M., Möller, H.E., 2019. Influence of the extracellular matrix on water mobility in subcortical gray matter. Magn. Reson. Med. 81: 1265–1279. https://doi.org/10.1002/mrm.27459.

31. Grad, J., Bryant, R.G., 1990. Nuclear magnetic cross-relaxation spectroscopy. J. Magn. Reson. 90: 1–8. https://doi.org/10.1002/mrm.1910170216.

32. Haase, A., Frahm, J., Matthaei, D., Hänicke, W., Merboldt, K.-D., 1986. FLASH imaging. Rapid NMR imaging using low flip-angle pulses. J. Magn. Reson. 67: 258–266. https://doi.org/10.1016/j.jmr.2011.09.021.

33. Helms, G., Dathe, H., Dechent, P., 2008. Quantitative FLASH MRI at 3T using a rational approximation of the Ernst equation. Magn. Reson. Med. 59: 667–672. https://doi.org/10.1002/mrm.21542.

34. Henkelman, R.M., Huang, X., Xiang, Q.-S., Stanisz, G.J., Swansosn, S.D., Bronskill, M.J., 1993. Quantitative interpretation of magnetization transfer. Magn. Reson. Med. 29: 759–766. https://doi.org/10.1002/mrm.1910290607.

35. Henkelman, R.M., Stanisz, G.J., Graham, S.J., 2001. Magnetization transfer in MRI: A review. NMR Biomed. 14: 57–64. https://doi.org/10.1002/nbm.683.

36. Hetzer, S., Mildner, T., Driesel, W., Weder, M., Möller, H.E., 2009. Shielded dual-loop resonator for arterial spin labeling at the neck. J. Magn. Reson. Imaging 29: 1414–1424. https://doi.org/10.1002/jmri.21803.

37. Hetzer, S., Mildner, T., Möller, H.E., 2011. A modified EPI sequence for high-resolution imaging at ultra-short echo time. Magn. Reson. Med. 65: 165–175. https://doi.org/10.1002/mrm.22610.

38. Horton, J.C., Hocking, D.R., 1997. Myelin patterns in V1 and V2 of normal and monocularly enucleated monkeys. Cereb. Cortex 7: 166–177. https://doi.org/10.1093/cercor/7.2.166.

39. Insko, E.K., Bolinger, L., 1993. Mapping of the radiofrequency field. J. Magn. Reson. A 103: 82–85. https://doi.org/10.1006/jmra.1993.1133.

40. Jeffery, G., Levitt, J.B., Cooper, H.M., 2008. Segregated hemispheric pathways through the optic chiasm distinguish primates from rodents. Neuroscience 157: 637–643. https://doi.org/10.1016/j.neuroscience.2008.09.021.

41. Jenkinson, M., Bannister, P., Brady, M., Smith, S., 2002. Improved optimization for the robust and accurate linear registration and motion correction of brain images. NeuroImage 17: 825–841. https://doi.org/10.1016/s1053-8119(02)91132-8.

42. Jenkinson, M., Beckmann, C.F., Behrens, T.E.J., Woolrich, M.W., Smith, S.M., 2012. FSL. NeuroImage 62: 782–790. https://doi.org/10.1016/j.neuroimage.2011.09.015.

43. Jenkinson, M., Smith, S., 2001. A global optimisation method for robust affine registration of brain images. Med. Image Anal. 5: 143–156. https://doi.org/10.1016/s1361-8415(01)00036-6.

44. Kiselev, V.G., Novikov, D.S., 2018. Transverse NMR relaxation in biological tissues. NeuroImage 182: 149–168. https://doi.org/10.1016/j.neuroimage.2018.06.002.

45. Koenig, S.H., 1991. Cholesterol of myelin is the determinant of gray-white contrast in MRI of brain. Magn. Reson. Med. 20: 285–291. https://doi.org/10.1002/mrm.1910200210.

46. Koenig, S.H., Brown III, R.D., Spiller, M., Lundbom, N., 1990. Relaxometry of brain: Why white matter appears bright in MRI. Magn. Reson. Med. 14: 482–495. https://doi.org/10.1002/mrm.1910140306.

47. Kozlowski, P., Raj, D., Liu, J., Lam, C., Yung, A.C., Tetzlaff, W., 2008. Characterizing white matter damage in rat spinal cord with quantitative MRI and histology. J. Neurotrauma 25: 653–676. https://doi.org/10.1089/neu.2007.0462.

48. Kucharczyk, W., Macdonald, P.M., Stanisz, G.J., Henkelman, R.M., 1994. Relaxivity and magnetization transfer of white matter lipids at MR imaging: Importance of cerebrosides and pH. Radiology 192: 521–529. https://doi.org/10.1148/radiology.192.2.8029426.

49. Labadie, C., Lee, J.-H., Rooney, W.D., Jarchow, S., Aubert-Frécon, M., Springer Jr., C.S., Möller, H.E., 2014. Myelin water mapping by spatially regularized longitudinal relaxographic imaging at high magnetic fields. Magn. Reson. Med. 71: 375–387. https://doi.org/10.1002/mrm.24670. Erratum. Magn. Reson. Med. 2015; 74: 1503. https://doi.org/10.1002/mrm.25974.

50. Larsen, M., Bjarkam, C.R., Stoltenberg, M., Sørensen, J.C., Danscher, G., 2003. An autometallographic technique for myelin staining in formaldehyde-fixed tissue. Histol. Histopathol. 18: 1125–1130. https://doi.org/10.14670/HH-18.1125.

51. Laule, C., Leung, E., Li, D.K.B., Traboulsee, A.L., Paty, D.W., MacKay, A.L., Moore, G.R.W., 2006. Myelin water imaging in multiple sclerosis: Quantitative correlations with histopathology. Mult. Scler. 12: 747–753. https://doi.org/10.1177/1352458506070928.

52. Laule, C., Vavasour, I.M., Kolind, S.H., Li, D.K.B., Traboulsee, T.L., Moore, G.R.W., MacKay, A.L., 2007. Magnetic resonance imaging of myelin. Neurotherapeutics 4: 460–484. https://doi.org/10.1016/j.nurt.2007.05.004.

53. Lenich, T., Pample, A., Mildner, T., Möller, H.E., 2019. A new approach to Z-spectrum acquisition: Prospective baseline enhancement (PROBE) for CEST/nuclear Overhauser effect. Magn. Reson. Med. 81: 2315–2329. https://doi.org/10.1002/mrm.27555.

54. Levesque, I.R., Pike., G.B., 2009. Characterizing healthy and diseased white matter using quantitative magnetization transfer and multicomponent T_2_ relaxometry: A unified view via a four-pool model. Magn. Reson. Med. 62: 1487–1496. https://doi.org/10.1002/mrm.22131.

55. Liu, J.V., Bock, N.A., Silva, A.C., 2011. Rapid high-resolution three-dimensional mapping of T1 and age-dependent variations in the non-human primate brain using magnetization-prepared rapid gradient-echo (MPRAGE) sequence. NeuroImage 56: 1154–1163. https://doi.org/10.1016/j.neuroimage.2011.02.075.

56. Lorio, S., Lutti, A., Kherif, F., Ruef, A., Dukart, J., Chowdhury, R., Frackowiak, R.S., Ashburner, J., Helms, G., Weiskopf, N., Draganski, B., 2014. Disentangling in vivo the effects of iron content and atrophy on the ageing human brain. NeuroImage: 280–289. https://doi.org/10.1016/j.neuroimage.2014.09.044.

57. Marques, J.P., Kober, T., Krueger, G., van der Zwaag, W., Van de Moortele, P.-F., Gruetter, R., 2010. MP2RAGE, a self bias-field corrected sequence for improved segmentation and T1-mapping at high field. NeuroImage 49: 1271–1281. https://doi.org/10.1016/j.neuroimage.2009.10.002.

58. McConnell, H.M., 1958. Reaction rates by nuclear magnetic resonance. J. Chem. Phys. 28: 430–431. https://doi.org/10.1063/1.1744152.

59. Meguro, R., Asano, A., Odagiri, S., Li, C., Shoumura, K., 2008. Cellular and subcellular localizations of nonheme ferric and ferrous iron in the rat brain: A light and electron microscopic study by the perfusion-Perls and -Turnbull methods. Arch. Histol. Cytol. 71: 205–222. https://doi.org/10.1679/aohc.71.205.

60. Mezer, A., Yeatman, J.D., Stikov, N., Kay, K.N., Cho, N.-J., Dougherty, R.F., Perry, M., Parvizi, J., Hua, L.H., Butts-Pauly, K., Wandell, B.A., 2013. Quantifying the local tissue volume and composition in individual brains with magnetic resonance imaging. Nat. Med. 19: 1667–1672. https://doi.org/10.1038/nm.3390.

61. Mispelter, J., Lupu, M., Briguet, A., 2006. NMR Probeheads for Biophysical and Biomedical Experiments. Theoretical Principles and Practical Guidelines. London: Imperial College Press; p. 280. https://doi.org/10.1142/P438.

62. Möller, H.E., Bossoni, L., Connor, J.R., Crichton, R.R., Does, M.D., Ward, R.J., Zecca, L., Zucca, F.A., Ronen, I., 2019. Iron, Myelin, and the brain: Neuroimaging meets neurobiology. Trends Neurosci. 42: 384–401. https://doi.org/10.1016/j.tins.2019.03.009.

63. Morris, C.M., Candy, J.M., Oakley, A.E., Bloxham, C.A., Edwardson, J.A., 1992. Histochemical distribution of non-haem iron in the human brain. Acta Anat. 144: 235–257. https://doi.org/10.1159/000147312.

64. Morrison, C., Stanisz, G., Henkelman, R.M., 1995. Modeling magnetization transfer for biological-like systems using a semi-solid pool with a super-lorentzian lineshape and dipolar reservoir. J. Magn. Reson. B 108: 103–113. https://doi.org/10.1006/jmrb.1995.1111.

65. Müller, D.K., Pampel, A., Möller, H.E., 2013. Matrix-algebra-based calculations of the time evolution of the binary spin-bath model for magnetization transfer. J. Magn. Reson. 230: 88–97. https://doi.org/10.1016/j.jmr.2013.01.013. Corrigendum. J. Magn. Reson. 2015; 261: 221. https://doi.org/10.1016/j.jmr.2015.11.001.

66. Müller, R., Pampel, A., Mildner, T., Marschner, H., Möller, H., 2013. A transceive RF coil for imaging tissue specimen at 3T based on PCB design. In Proceedings of the 21st Annual Meeting of ISMRM, Salt Lake City, UT, USA, p. 4366.

67. Newman, J.D., Kenkel, W.M., Aronoff, E.C., Bock, N.A., Zametkin, M.R., Silva, A.C., 2009. A combined histological and MRI brain atlas of the common marmoset monkey, Callithrix jacchus. Brain Res. Rev. 62: 1–18. https://doi.org/10.1016/j.brainresrev.2009.09.001.

68. Nieuwenhuys R., 2012. The myeloarchitectonic studies on the human cerebral cortex of the Vogt-Vogt school, and their significance for the interpretation of functional neuroimaging data. Brain Struct. Funct. 218: 303–352. https://doi.org/10.1007/s00429-012-0460-z.

69. Norton, W.T., Cammer, W., 1984. Isolation and characterization of myelin. In: Morell, P. (ed.). Myelin. Second Edition. New York, NY: Springer Science+Business Media; p. 147–196.

70. Norton, W.T., Poduslo, S.R., 1973. Myelination in rat brain: Changes in myelin composition during brain maturation. J. Neurochem. 21: 759–773. https://doi.org/10.1111/j.1471-4159.1973.tb07520.x.

71. O’Brian, J.S., Sampson, E.L., 1965. Lipid composition of the normal human brain: Gray matter, white matter, and myelin. J. Lipid Res. 6: 537–544. https://doi.org/10.1016/S0022-2275(20)39619-X.

72. Palomero-Gallagher, N., Zilles, K., 2019. Cortical layers: Cyto-, myelo, receptor- and synaptic architecture in human cortical areas. NeuroImage 197: 716–741. https://doi.org/10.1016/j.neuroimage.2017.08.035.

73. Pampel, A., Müller, D.K., Anwander, A., Marschner, H., Möller, H.E., 2015. Orientation dependence of magnetization transfer parameters in human white matter. NeuroImage 114: 136–146. https://doi.org/10.1016/j.neuroimage.2015.03.068.

74. Paxinos, G., Watson, C., Petrides, M., Rosa, M., Hironobu, T., 2012. The Marmoset Brain in Stereotaxic Coordinates. London: Academic Press, 282 p.

75. Pistorio, A.L., Hendry, S.H., Wang, X., 2006. A modified technique for high-resolution staining of myelin. J. Neurosci. Methods 153: 135–146. https://doi.org/10.1016/j.jneumeth.2005.10.014.

76. Portnoy, S., Stanisz, G.J., 2007. Modeling pulsed magnetization transfer. Magn. Reson. Med. 58: 144–155. https://doi.org/10.1002/mrm.21244.

77. Reinert, A., Morawski, M., Seeger, J., Arendt, T., Reinert, T., 2019. Iron concentrations in neurons and glial cells with estimates on ferritin concentrations. BMC Neurosci. 20: 25. https://doi.org/10.1186/s12868-019-0507-7.

78. Salustri, C., 1996. Lack of magnetization transfer from the ferritin molecule. J. Magn. Reson. B 111: 171–173. https://doi.org/10.1006/jmrb.1996.0076.

79. Schmierer, K., Tozer, D.J., Scaravilli, F., Altmann, D.R., Barker, G.J., Tofts, P.S., Miller, D.H., 2007. Quantitative magnetization transfer imaging in postmortem multiple sclerosis brain. J. Magn. Reson. Imaging 26: 41–51. https://doi.org/10.1002/jmri.20984.

80. Sled, J.G., 2018. Modelling and interpretation of magnetization transfer imaging in the brain. NeuroImage 182: 128–135. https://doi.org/10.1016/j.neuroimage.2017.11.065.

81. Sled, J.G., Levesque, I., Santos, A.C., Francis, S.J., Narayanan, S., Brass, S.D., Arnold, D.L., Pike, G.B., 2004. Regional variations in normal brain shown by quantitative magnetization transfer imaging. Magn. Reson. Med. 51: 299–303. https://doi.org/10.1002/mrm.10701.

82. Smith, S.A., Bulte, J.W.M., van Zijl, P.C.M., 2009. Direct saturation MRI: Theory and application to imaging brain iron. Magn. Reson. Med. 62: 384–393. https://doi.org/10.1016/j.neuroimage.2017.11.065.

83. Stanisz, G.J., Kecojevic, A., Bronskill, M.J., Henkelman, R.M., 1999. Characterizing white matter with magnetization transfer and ??. Magn. Reson. Med. 42: 1128–1136. https://doi.org/10.1002/mrm.21980.

84. Stikov, N., Campbell, J.S.W., Stroh, T., Lavelée, M., Frey, S., Novek, J., Nuara, S., Ho, M.-K., Bedell, B.J., Dougherty, R.F., Leppert, I.R., Boudreau, M., Narayanan, S., Duval, T., Cohen-Adad, J., Picard, P.-A., Gasecka, A., Côté, D., Pike, G.B., 2015. In vivo histology of the myelin g-ratio with magnetic resonance imaging. NeuroImage 118: 397–405. https://doi.org/10.1016/j.neuroimage.2015.05.023.

85. Stüber, C., Morawski, M., Schäfer, A., Labadie, C., Wähnert, M., Leuze, C., Streicher, M., Barapatre, N., Reimann, K., Geyer, S., Spemann, D., Turner, R., 2014. Myelin and iron concentration in the human brain: A quantitative study of MRI contrast. NeuroImage 93: 95–106. https://doi.org/10.1016/j.neuroimage.2014.02.026.

86. Tomassy, G.S., Berger, D.R., Chen, H.-H., Kasthuri, N., Hayworth, K.J., Vercelly, A., Seung, H.S., Lichtman, J.W., Arlotta, P., 2014. Distinct profiles of myelin distribution along single axons of pyramidal neurons in the neocortex. Science 344: 319–324. https://doi.org/10.1126/science.1249766.

87. Trujillo, P., Summers, P.E., Ferrari, E., Zucca, F.A., Sturini, M., Mainardi, L.T., Cerutti, S., Smith, A.K., Smith, S.A., Zecca, L., Costa, A., 2017. Contrast mechanisms associated with neuromelanin-MRI. Magn. Reson. Med. 78: 1790–1800. https://doi.org/10.1002/mrm.26584.

88. Tyler, D.J., Gowland, P.A., 2005. Rapid quantitation of magnetization transfer using pulsed off-resonance irradiation and echo planar imaging. Magn. Reson. Med. 53: 103–109. https://doi.org/10.1002/mrm.20323.

89. Uchihara, T., 2007. Silver diagnosis in neuropathology: Principles, practice and revised interpretation. Acta Neuropathol. 113: 483–499. https://doi.org/10.1007/s00401-007-0200-2.

90. Vavasour, I.M., Whittall, K.P., MacKay, A.L., Li, D.K.B, Vorobeychik, G., Paty, D.W., 1998. A comparison between magnetization transfer ratios and myelin water percentages in normals and multiple sclerosis patients. Magn. Reson. Med. 40: 763–768. https://doi.org/10.1002/mrm.1910400518.

91. Watanabe, O., West, C.R., Bremer, A., 1977. Experimental regional cerebral ischemia in the middle cerebral artery territory in primates. Part 2: Effects on brain water and electrolytes in the early phase of MCA stroke. Stroke 8: 71–76. https://doi.org/10.1161/01.str.8.1.71.

92. Weiss, M., Alkemade, A., Keuken, M.C., Müller-Axt, C., Geyer, S., Turner, R., Forstmann, B.U., 2015. Spatial normalization of ultrahigh resolution 7 T magnetic resonance imaging data of the postmortem human subthalamic nucleus: A multistage approach. Brain Struct. Funct. 220: 1695–1703. https://doi.org/10.1007/s00429-014-0754-4.

93. Wennerström, H., 1973. Proton nuclear magnetic resonance lineshapes in lamellar liquid crystals. Chem. Phys. Lett. 18: 41–44. https://doi.org/10.1016/0009-2614(73)80333-1.

94. Wolff, S.D., Balaban, R.S., 1989. Magnetization transfer contrast (MTC) and tissue water proton relaxation in vivo. Magn. Reson. Med. 10: 135–144. https://doi.org/10.1002/mrm.1910100113.

95. Yuasa, S., Nakamura, K., Kohsaka, S., 2010. Stereotaxic Atlas of the Marmoset Brain. With Immunohistochemical Architecture and MR Images. Tokyo: National Institute of Neuroscience.

