## Supplementary Information for "High-Resolution Magnetization-Transfer Imaging of *Post-Mortem* Marmoset Brain: Comparisons with Relaxometry and Histology"

---

### Abbreviations

2D = two-dimensional; 3D = three-dimensional; 3V = 3rd ventricle; A24 = area 24; ac = anterior commissure; Am = amygdaloid nuclei; BSB = binary spin bath; c = central part; cc = corpus callosum; Cd = caudate nucleus; Cl = claustrum; cr = corona radiata; ec = external capsule; EN = endopiriform nuclei; ex = extreme capsule; FLASH = fast low-angle shot; FOV = field of view; GM = gray matter; ic = internal capsule; Ins = insula; IOD = integrated optical density; l = lateral part; lf = lateral fissure; LV = lateral ventricle; M1 = primary motor cortex; MBP = myelin basic protein; ME = multi echo; MP2RAGE = magnetization-prepared 2 rapid gradient echoes; MR = magnetic resonance; MRI = magnetic resonance imaging; MT = magnetization transfer; och = optic chiasm; PaC = parietal cortex; PBS = phosphate-buffered saline; PCB = printed circuit board; Pu = putamen; qMTI = quantitative magnetization transfer imaging; RF = radiofrequency; ROI = region of interest; S1 = primary somatosensory cortex; SNR = signal-to-noise ratio; V1 = primary visual cortex; VP = ventral pallidum; WM = white matter.

### Mathematical Symbols

$A$: absorbance;
$a$: normalized absorbance;
$^{a}$, $^{b}$: superscripts denoting the free and the semisolid pool, respectively;
$_{\mathrm{app}}$: subscript denoting an apparent quantity;
$\mathbf{B}_0$: static magnetic field vector;
$B_0$: static magnetic field amplitude;
$B_1^+$: RF transmit magnetic field amplitude;
$C$: capacitance;
$C_m$: matching capacitance;
$C_t$: tuning capacitance;
$c_1, c_0, c_0'$: linear coefficients in multivariate regression analysis;
$E_1$: exponential longitudinal relaxation factor;
$F$: ratio of mean squared error and mean squared residuals;
$\mathcal{F}$: pool-size ratio $M_0^b/M_0^a$;
$_{\mathrm{Fe}}$: subscript denoting Perls' stain for ferric iron ($Fe^{3+}$);
$f$: fraction of the total tissue mass;
$f_{\mathrm{max}}^m$: maximum myelin fraction of the total tissue mass;
$g^b$: absorption lineshape function of the semisolid pool;
$I$: transmitted intensity;
$I_0$: incident intensity;
$i$: position index;
$^{iew}$, $^{mw}$: superscripts denoting intra-/extracellular and myelin water, respectively;
$M_0$: equilibrium magnetization;
$M_z$: longitudinal magnetization;
$_{\mathrm{MBP}}$: subscript denoting MBP immunostaining;
MTR magnetization transfer ratio;
MWF myelin water fraction;
$^{m}$, $^{nm}$: superscripts denoting myelin and non-myelin dry matter, respectively;
$_{\mathrm{my}}$: subscript denoting Gallyas' silver stain for myelin;
$N_{\mathrm{av}}$: number of averages;
$n$: number of voxels;
$p$: error probability;
$Q$: quality factor;
$R$: exchange rate constant;
$R_1$: longitudinal relaxation rate;
$R_1^{\mathrm{obs}}$: observed longitudinal relaxation rate;
$R_2$: transverse relaxation rate;
$R_2^*$: effective transverse relaxation rate;
$R_{\mathrm{RF}}$: RF saturation rate;
RSME: root mean-squared error;
$r$: Pearson correlation coefficient;

$r^2_{\mathrm{adj}}$: adjusted coefficient of determination;
$S$: signal voltage;
$S_0$: signal voltage generated by a 90° pulse to the fully relaxed magnetization;
SD: standard deviation;
$T_1$: longitudinal relaxation time;
$T_1^{\mathrm{obs}}$: observed longitudinal relaxation time;
$T_2$: transverse relaxation time;
$T_2^*$: effective transverse relaxation time;
TE: echo time;
ΔTE: inter-echo time;
TI: inversion time;
TR: repetition time;
$x, y, z$: Cartesian coordinates;
$y(x_1, x_2, \dots)$: function of the variables $x_1$, $x_2$, …;
$\alpha$: imaging pulse flip angle;
$\vartheta$: polar angle (in the magnet coordinate system);
$\epsilon_1, \epsilon_0$: linear coefficients describing the correlation of $a_{\mathrm{my}}$ and $a_{\mathrm{Fe}}$;
$\sigma$: scaling factor;
$\tau_p$: pulse duration;
$\varphi$: azimuth angle (in the magnet coordinate system);
$\Omega$: off-resonance angular frequency;
$\omega_1$: RF pulse amplitude (in rad/s);
$\omega_{1,\mathrm{max}}$: RF pulse peak amplitude (in rad/s);
$\bar{x}$: mean value of the quantity $x$.

**Supplementary Table S1.** Pearson correlation coefficients, $r$, for pairwise comparisons of histology results (normalized IOD of Gallyas' myelin stain, MBP immunostain, and Perls' iron stain), MT parameters (pool-size ratio $\mathcal{F} = M_0^b/M_0^a$ and exchange rate), and relaxation rates $R_1^{\text{obs}}$ (at 3 T and 7 T) and $R_2^*$. The upper-right (gray background) and lower-left (white background) parts of the correlation diagram show results from separate analyses in all GM ($n$ = 4,964) and all WM voxels ($n$ = 1,538; including optic chiasm), respectively, which were performed without outlier removal. Significant correlations (after Bonferroni correction) are indicated in bold, numbers in brackets indicate insignificant correlations.

| $r$ | $a_{\text{my}}$ | $a_{\text{MBP}}$ | $a_{\text{Fe}}$ | $\mathcal{F}$ | $RM_0^a$ | $R_{1,3\text{T}}^{\text{obs}}$ | $R_{1,7\text{T}}^{\text{obs}}$ | $R_2^*$ | |
|---|---|---|---|---|---|---|---|---|---|
| $a_{\text{my}}$ | | **0.894****** | **0.568****** | **0.531****** | **0.454****** | **0.633****** | **0.675****** | **0.474****** | GM |
| $a_{\text{MBP}}$ | *(0.041)* | | **0.626****** | **0.629****** | **0.438****** | **0.672****** | **0.711****** | **0.573****** | |
| $a_{\text{Fe}}$ | **0.411****** | *( 0.056)* | | **0.607****** | **0.272****** | **0.614****** | **0.630****** | **0.669****** | |
| $\mathcal{F}$ | **0.573****** | **–0.161****** | **0.537****** | | **0.254****** | **0.714****** | **0.704****** | **0.618****** | |
| $RM_0^a$ | **0.156****** | **0.348****** | **0.165****** | *(0.028)* | | **0.484****** | **0.553****** | **0.376****** | |
| $R_{1,3\text{T}}^{\text{obs}}$ | **0.571****** | *( 0.076)* | **0.647****** | **0.730****** | **0.371****** | | **0.871****** | **0.655****** | |
| $R_{1,7\text{T}}^{\text{obs}}$ | **0.619****** | *( 0.022)* | **0.645****** | **0.734****** | **0.327****** | **0.872****** | | **0.739****** | |
| $R_2^*$ | **0.522****** | *(–0.027)* | **0.637****** | **0.661****** | **0.219****** | **0.748****** | **0.824****** | | |
| | WM | | | | | | | | |

*** $p < 0.001$ (corrected).

**Supplementary Table S2.** $F$-statistic results including the root mean-squared error (RMSE), adjusted coefficient of determination ($r_{\text{adj}}^2$), $F$-value and $p$-value (Bonferroni-corrected) for comparisons of bivariate linear regression models $y(x_\alpha, x_\beta) = c_1^\alpha x_\alpha + c_1^\beta x_\beta + c_0$ (see Table 4) to a constant model.

| | | | Whole slice | | | GM mask | | | WM mask | | |
|---|---|---|---|---|---|---|---|---|---|---|---|
| $y(x_\alpha, x_\beta)$ | $x_\alpha$ | $x_\beta$ | RMSE | $r_{\text{adj}}^2$ | $F$ | RMSE | $r_{\text{adj}}^2$ | $F$ | RMSE | $r_{\text{adj}}^2$ | $F$ |
| $R_{1,3\text{T}}^{\text{obs}}/\text{s}^{-1}$ | $a_{\text{my}}$ | $a_{\text{Fe}}$ | 0.178 | 0.713 | 7,330*** | 0.146 | 0.496 | 2,440*** | 0.247 | 0.572 | 613*** |
| $R_{1,3\text{T}}^{\text{obs}}/\text{s}^{-1}$ | $\mathcal{F}$ | $a_{\text{Fe}}$ | 0.149 | 0.799 | 11,700*** | 0.136 | 0.561 | 3,170*** | 0.194 | 0.735 | 1,270*** |
| $R_{1,7\text{T}}^{\text{obs}}/\text{s}^{-1}$ | $a_{\text{my}}$ | $a_{\text{Fe}}$ | 0.164 | 0.759 | 9,250*** | 0.14 | 0.546 | 2,980*** | 0.201 | 0.576 | 626*** |
| $R_{1,7\text{T}}^{\text{obs}}/\text{s}^{-1}$ | $\mathcal{F}$ | $a_{\text{Fe}}$ | 0.145 | 0.812 | 12,700*** | 0.138 | 0.56 | 3,160*** | 0.173 | 0.687 | 1,010*** |
| $R_2^*/\text{s}^{-1}$ | $a_{\text{my}}$ | $a_{\text{Fe}}$ | 5.73 | 0.687 | 6,470*** | 4.76 | 0.461 | 2,120*** | 7.52 | 0.442 | 365*** |
| $R_2^*/\text{s}^{-1}$ | $\mathcal{F}$ | $a_{\text{Fe}}$ | 5.02 | 0.76 | 9,310*** | 4.49 | 0.519 | 2,680*** | 7.08 | 0.506 | 472*** |
| $\mathcal{F}$ | $a_{\text{my}}$ | $a_{\text{Fe}}$ | 0.0278 | 0.664 | 5,830*** | 0.0156 | 0.419 | 1,790*** | 0.039 | 0.354 | 253*** |
| $RM_0^a/\text{s}^{-1}$ | $a_{\text{my}}$ | $a_{\text{Fe}}$ | 2.05 | 0.209 | 780*** | 1.96 | 0.206 | 645*** | 2.3 | 0.128 | 69*** |

*** $p < 0.001$ (corrected).
